# Identification and characterization of a human antibody profile inversely associated with adverse cardiovascular events

**DOI:** 10.64898/2026.09.01.748614

**Authors:** David Henson, Katherine L. Thompson, Ayman Samman Tahhan, Ryan Marion, Tabarak Azawi, Peng Yeh, Gregory S. Hawk, Andrew DeFilippis, Arshed Ali Quyyumi, Vincent J. Venditto

## Abstract

**Objective:** The immune response is linked to the progression of atherosclerotic cardiovascular disease (CVD). We sought to characterize a predictive cardiovascular antibody biomarker using two independent cohorts.

**Approach and Results:** A human IgG profile serendipitously identified by ELISA was quantified in a well characterized cohort of 359 patients with a history of coronary artery disease (CAD) who experienced a total of 71 incident adverse CVD events (death, myocardial infarction, and stroke) over a median 4.1-year follow-up. Using Cox proportional hazard regression analysis, low biomarker levels were an independent predictor of adverse cardiovascular outcomes after adjustment for age, sex, diabetes mellitus, estimated glomerular filtration rate, presence of obstructive CAD, heart failure, total cholesterol, and high-density lipoprotein (HDL) cholesterol (adjusted hazard ratio of 1.90 [95% CI: 1.03–3.49; *p*=0.038] between lowest and highest tertiles). Validation was then performed in a larger secondary cohort using 4356 baseline samples from the Multi-Ethnic Study of Atherosclerosis (MESA), a prospective study of cardiovascular outcomes resulting in adjusted odds ratio of 50.5 per unit decrease in log-transformed biomarker [95% CI: 2.4–2773.1; *p*=0.030] over one year by logistic regression analysis.

**Conclusions:** Low levels of human IgG antibodies targeting *Bovidae* IgG are independently associated with increased incidence of CVD events in patients with or without a history of CAD, indicating the potential clinical predictive power of this antibody profile as an inverse biomarker for CVD.

## INTRODUCTION

The inflammatory immune response is a significant driver of cardiovascular disease (CVD), resulting in increased rates of morbidity and mortality in patients around the world.^1, 2^ The role of the immune response and the elevated risk of major adverse cardiovascular events in subjects with autoimmunity and other chronic inflammatory conditions^3^ have led to the identification of immunologic biomarkers with the potential to improve risk stratification of patients with CVD.^4^ These biomarkers include general inflammatory markers,^5, 6^ specific antibodies with known targets,^7^ and immune complexes (ICs) containing immunoglobulin and antigen.^8, 9^ Assay development and validation are critical to quantify biomarkers to assess correlations with disease outcomes.

Rational assay development to quantify predetermined analytes provides a clear route to biomarker correlation with disease. However, assay interference derived from unintended target detection in human samples remains a confounding factor, including those used for clinical care.^10^ One form of interference occurs when human antibodies, present in clinical samples, bind to the assay reagents to enhance or suppress assay values.^11, 12^ Such antibodies are described as either heterophilic antibodies, classified as natural antibodies with low affinity, or human anti-animal antibodies (HAAA) which bind with high affinity to their cognate antigen.^12^ HAAA can be induced through iatrogenic means using animal-derived pharmaceuticals resulting in antibodies targeting mouse, rat, rabbit, pig, and other species.^11^ Non-iatrogenic activities, such as animal husbandry^13^ and dietary consumption,^14^ also result in increased levels of HAAA leading to variability in the seroprevalence and specificity of HAAA across populations.

Alternatively, antibodies to animal products can result from molecular mimicry in which an immune response is induced toward an unrelated antigen with sequence similarity causing cross-reactivity.^15^ Regardless of the stimuli for induction, HAAA are linked with interference in clinical assays resulting in misdiagnosis and unnecessary treatment,^11, 16–18^ but their role as biomarkers of disease have not been explored.

In this study, we sought to characterize an antibody biomarker that inversely correlates with cardiovascular outcomes, whose target antigen was incorrectly assigned in a previous report.^19^ We originally hypothesized that immune complexes between apolipoprotein A-I and IgG (ApoA-I/IgG ICs) may serve as a biomarker for CVD events, based on evidence of anti-ApoA-I antibodies in human subjects and mouse samples.^20–30^ To test this hypothesis, we developed an ELISA to measure ApoA-I/IgG ICs and utilized this assay to analyze samples from patients with a history of coronary artery disease (CAD) to investigate their association with CVD events. Although a correlation with cardiovascular outcomes was observed, continued characterization indicated that the observed antibody profile is not an IC as previously reported,^19^ but a human antibody reactive with goat-IgG used in the assay. Herein, we describe the development of the assay, associated outcomes in a cohort of patients with CAD, characterization of the biomarker, and confirmation of the cardiovascular outcomes using baseline samples from the Multi-Ethnic Study of Atherosclerosis (MESA) as a prospective population and an independent cohort for biomarker validation.

## MATERIALS AND METHODS

### Human samples

#### Ethics Statement

The original MESA study was conducted in accordance with the Declaration of Jelsinki, and its protocols were approved by the Institutional Review Boards of all collaborating data collection sites. Written informed consent was obtained from all individual participants included in the study. For the secondary analysis, ethical approval was obtained from the University of Kentucky Institutional Review Board (protocol #: 52074).

#### Patients With CAD

Samples were obtained from 359 patients with CAD enrolled in the Emory Cardiovascular Biobank, a prospective cohort of 20- to 90-year-old patients who underwent elective or emergent cardiac catheterization.^31^ Arterial blood was collected into EDTA tubes and plasma isolated from whole blood prior to storage until use at −80 °C. All subjects were provided informed consent. Hypertension, hypercholesterolemia, and diabetes mellitus were defined according to the Joint National Committee, Adult Treatment Panel III, and American Diabetes Association criteria, respectively.^32–34^ Additional patient information is provided in the Supporting Materials.

### Blood Donor Subjects

De-identified human blood samples were obtained from the Kentucky Blood Center (KBC) for analysis. During the blood donation process at the KBC, additional blood samples are collected in EDTA tubes and stored at 4 °C for 7 days for follow-up testing as needed. Remaining samples after testing was completed were provided to our laboratory. Upon receiving samples, plasma was separated from the whole blood via centrifugation and stored at −80 °C until assayed. *Blood donor sample characteristics*: A subset of 112 KBC blood donor plasma samples were utilized for experiments described. All 112 samples exhibit an absorbance at least three standard deviations above background indicating the presence of biomarker. The 15 samples used for detailed characterization (protein G purification, and western blot) were selected based on the following criteria: Nine samples with biomarker values near the top of the cohort, three samples at the 66^th^ percentile and three samples with low values. For Figures S1, S2, and S6 alternative blood donor samples were utilized based on available sample volume.

### MESA Plasma Samples

The Multiethnic Study of Atherosclerosis (MESA) enrolled 6,814 participants (3,601 women, 3,213 men) with no known clinical atherosclerotic CVD of four racial/ethnic backgrounds (White, Chinese, Black, and Hispanic/Latino), aged 45-84 years as described in detail previously.^35^ An ancillary study conducted by our group measured biomarker levels at baseline in 4356 MESA participants. Samples were analyzed in numerical order using the Central Blood Analysis Laboratory (CBAL) number. The population tested reflects a portion of the full MESA cohort as only samples tested using reagents from matched lot numbers were included to minimize confounding variability when using assay reagents from different lots.

### Biochemical Analyses

#### Biomarker ELISA

Biomarker levels were quantified by ELISA using streptavidin coated plates incubated consecutively with biotinylated capture antibodies, plasma, and horseradish peroxidase conjugated secondary antibodies. Samples were evaluated in duplicate and analyzed for coefficient of variation (CV) to confirm low replicate variability. Samples with CV > 10% were reanalyzed. ELISA validation assays are shown in **Figure S1 and S2**. Competition ELISAs were performed using increasing concentrations of free goat IgG incubated with the biomarker plasma prior to ELISA assay. Complete details and lot numbers of the assays are provided in the Supporting Information.

### Western Blot

Biomarker interaction with goat-IgG were determined by western blot by using alternating lanes with 7 goat IgG samples and 8 ladders per gel. After electrophoresis and membrane transfer, membranes were cut such that one goat IgG lane and at least one ladder were visible on each membrane. The membranes were blotted with different human plasma samples and visualized with enhanced chemiluminescence (ECL). Complete details and lot numbers of the assays are provided in the Supporting Information.

### Bio-Layer Interferometry (BLITz) Assay

Binding kinetics of the biomarker were measured by BLITz assay using both goat-IgG and human-IgG immobilized on the sensor. Biomarker association was measured by placing the probe in human plasma samples, while dissociation was measured using assay buffer. Complete details and lot numbers of the assays are provided in the Supporting Information.

### Statistical Analysis

Statistical analysis was completed using Graph Pad Prism, SPSS, and R software. Spearman and Pearson correlations were used to evaluate relationships with clinical characteristics from patients with CAD. Subject characteristics were reported as descriptive statistics with means and standard deviations. Kaplan-Meier curves and Cox regression models were used to examine the association between biomarker levels and the composite outcome of death/MI/stroke. Cox regression models were adjusted for age, sex, diabetes mellitus, estimated glomerular filtration rate, presence of obstructive CAD, heart failure, total cholesterol, and HDL cholesterol. Characteristics incorporated in multivariable analyses included variables that correlated in bivariate analysis with the outcome studied. For group comparison, data was tested for normality using the Shapiro Wilk test and for equality of variance using the Brown Forsythe test. When appropriate, based on these results, samples were compared using Repeated Measures ANOVA with Bonferroni’s multiple comparisons. If ANOVA was not appropriate, samples were compared with Friedman’s test with Dunn’s multiple comparisons.

For the MESA cohort analysis, MESA data entry codes corresponding to clinical variables are described in detail in the Supporting Material. Cox regression models and logistic regression models were fit to the primary endpoint (Cardiovascular Disease (CVD), All [cvda]) with the quantitative log-transformed biomarker at each timepoint (1-, 2-, 3-, 4-, or 18.5-year post blood draw) as the primary explanatory variable of interest. Additional logistic regression models were also used to adjust for covariates across three models (Model 1: based on variables used in primary cohort; Model 2: based on variables used in PREVENT risk calculator^36^; Model 3: based on variables used in the Pooled Cohort Equation (PCE) risk calculator^37^). Wald’s tests were used to determine significance of the biomarker effect in each case. Both unadjusted and adjusted logistic regression models were fit individually to each diagnosis as a secondary endpoint (coronary heart disease all [chda], cardiovascular death [dthtype], all-cause mortality [dth], myocardial infarct [mi], stroke [strk]). All tests were considered statistically significant when *p*<0.05.

## RESULTS

### Biomarker quantification

To evaluate ApoA-I/IgG ICs as a potential biomarker for CVD progression we performed the ELISA in a cohort of 359 patients with a history of CAD. Demographic and clinical characteristics are shown in **Table S1**; mean age was 64 years, 65% males, 24% Black. Of the cohort, 88% had a history of hypertension, 77% had hyperlipidemia, and 34% had diabetes mellitus. Approximately 99% of the 359 samples exhibit detectable levels in the biomarker assay using a cutoff three standard deviations above background (Mean: 0.27 ± 0.21; IQR: 0.19-0.28) (**Figure 1A**). No associations were observed between the response in the ELISA and either the demographic characteristics or cardiovascular risk factors (**Table S2**).

**Figure 1.**
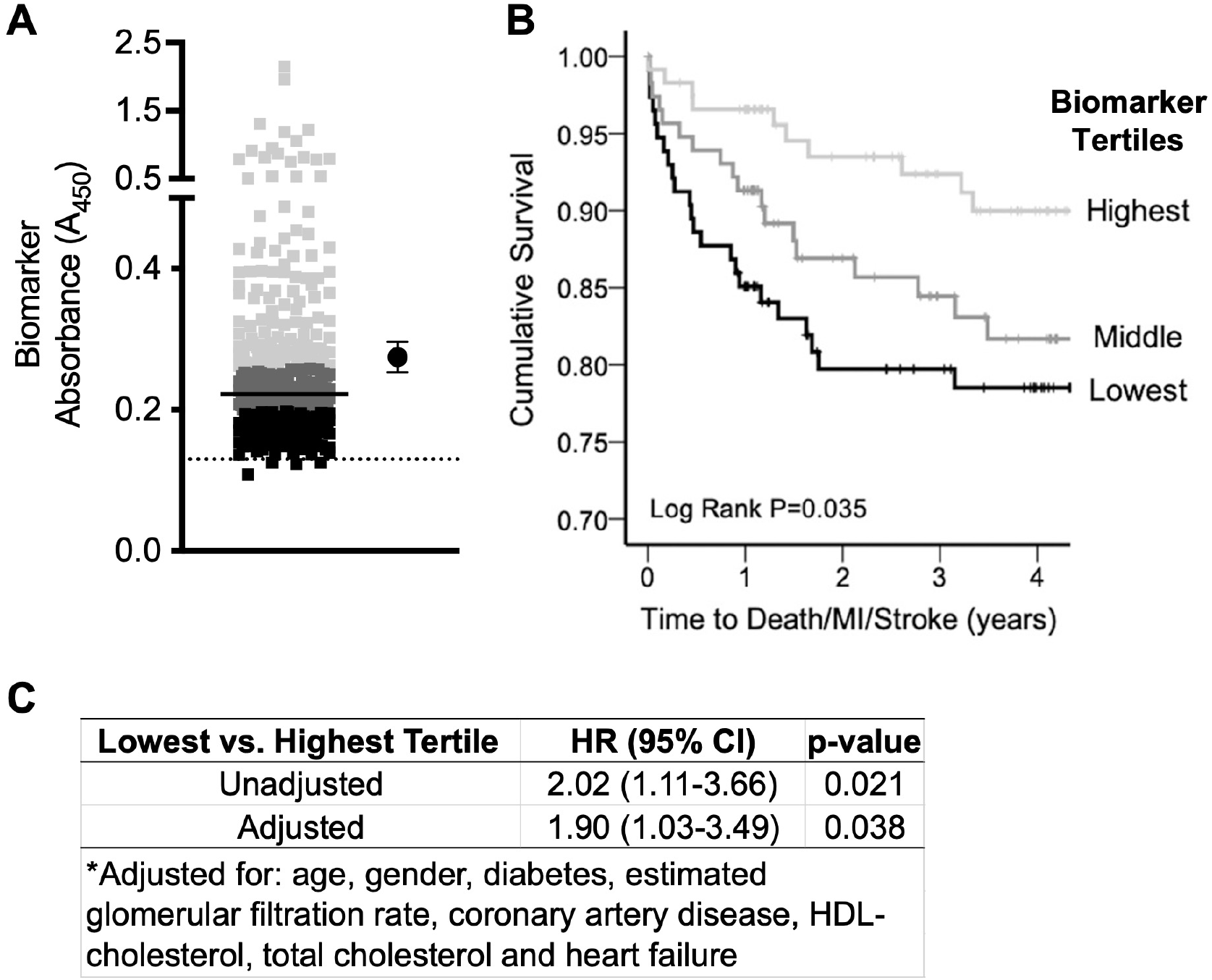
Cox proportional regression analysis of patients with CAD based on biomarker levels. **A**. Biomarker levels were evaluated by ELISA at a plasma dilution of 1:200 in 359 patients with CAD after undergoing emergent or elective cardiac catheterization. Absorbance values are plotted, with median denoted by line, tertiles are color coded to indicate high (light gray), middle (dark gray), and low (black). The mean value and 95% CI are plotted to the right. The dashed line denotes the limit of detection, defined as 3 SD above plasma-free wells. **B**. Kaplan-Meier plot of major adverse cardiovascular events over a median of 4.1 years (interquartile range, 1.2–6.0 y) of follow-up in patients with CAD. Patients are divided into tertiles by absorbance in the biomarker assay as denoted in **A**. **C**. Proportional regression models used to determine hazard ratios and CIs based on traditional markers of cardiovascular disease. HDL indicates high-density lipoprotein; and MI, myocardial infarction.

Subsequently, patients were divided into tertiles based on biomarker level to determine the association between the biomarker and CVD events. There were 71 adverse CVD events in this cohort defined as all-cause mortality, non-fatal MI and non-fatal stroke during a median 4.1-year (interquartile range, 1.2–6.0 years) follow-up. As shown in **Figure 1B**, the rate of adverse CVD events was higher in those with lower biomarker (unadjusted hazard ratio (HR) 2.02 [95% CI, 1.11–3.66; *p*=0.021] between the lowest and highest tertiles; **Figure 1C**). After Cox proportional hazards regression analysis adjusted for age, sex, diabetes mellitus, hypertension, estimated glomerular filtration rate, presence of obstructive CAD, heart failure, HDL cholesterol, and total cholesterol, low biomarker levels remained an independent predictor of incident adverse events (HR 1.90 [95% CI, 1.03–3.49; *p*=0.038] between the lowest and highest tertiles; **Figure 1C**).

### Biomarker Characterization

For further biomarker characterization, we modified the assay using capture antibodies targeting both self (Apo-B)^8, 9^ and non-self (green fluorescent protein (GFP)) antigens using 112 anonymized blood samples from the Kentucky Blood Center (**Figure 2A, B**). Notably, all three antibodies are derived from goat. Correlation analysis indicates similar biomarker levels between antibodies when compared to anti-ApoA-I (anti-ApoB: slope = 0.97 ± 0.13 (95% CI), R^2^ = 0.65; anti-GFP: slope = 1.02 ± 0.08, R^2^ = 0.86) (**Figure S3**). Importantly, the biomarker level is not due to non-specific binding in uncoated plates lacking capture antibody (slope = 0.20 ± 0.07; R^2^ = 0.21) (**Figure S3**), or due to non-specific binding with the secondary antibody (**Figure S4**). These data suggest that the biomarker is not an IC, but likely an artifact of assay interference related to the source of capture antibody.

**Figure 2.**
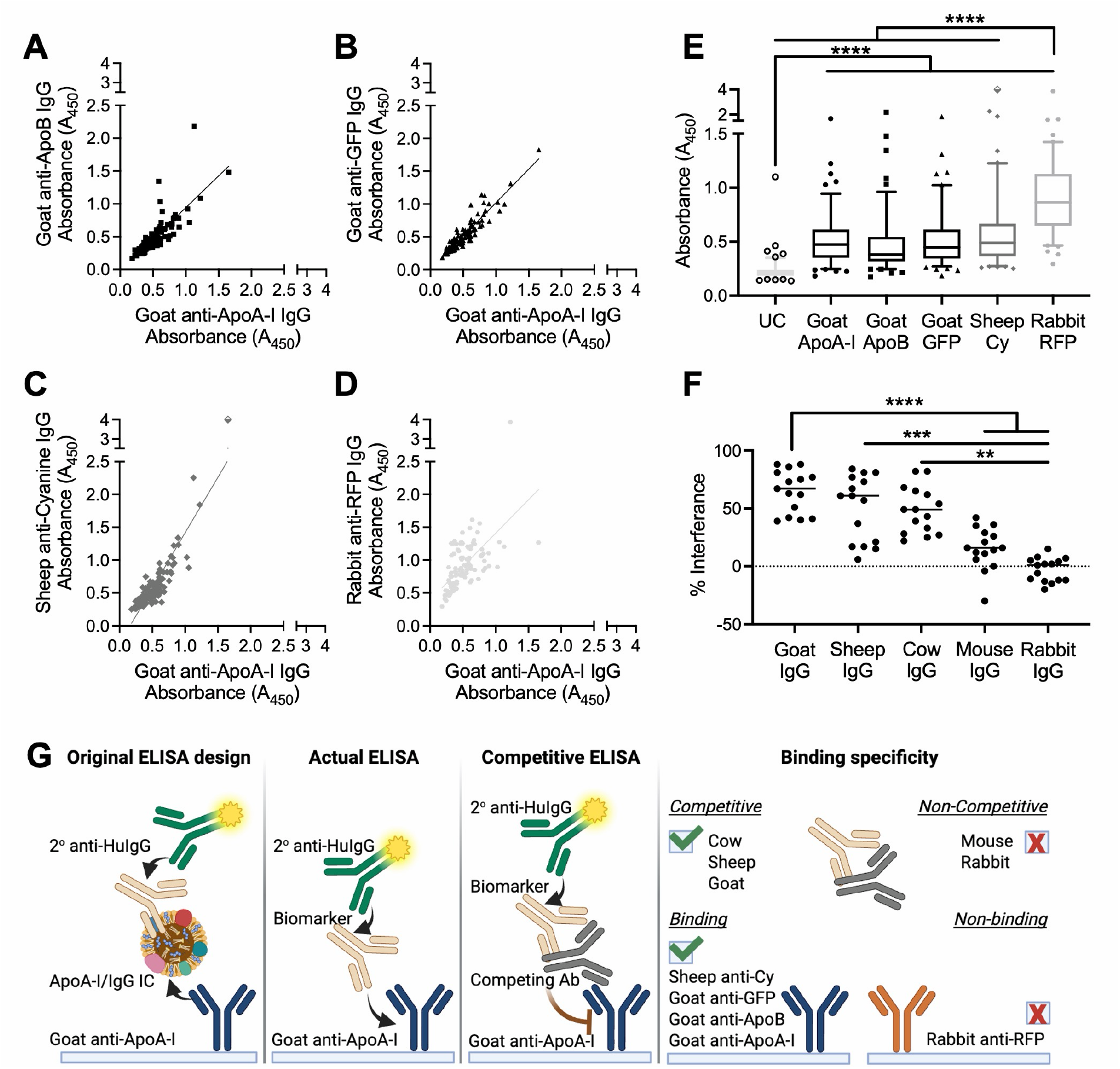
Effects of capture antibody used in the biomarker ELISA. **A**-**D**. Absorbance using goat anti-ApoA-I IgG as capture antibody in comparison to: **A**. Goat anti-ApoB IgG, **B**. Goat anti-GFP IgG, **C** Sheep anti-cyanine IgG, and **D**. Rabbit anti-RFP IgG as capture antibodies. All assays are performed with 112 plasma samples at a dilution of 1:200. **E**. Box and whisker plot of absorbance values from the 112 samples as shown in panels. **F**. Percent interference with pre-incubation of plasma with 500x IgG from corresponding species. Statistics in **E** and **F** performed using Friedman’s Test with Dunn’s multiple comparisons. **G** Summary of original designed ELISA, proposed actual ELISA based on antibody specificity, competitive ELISA, and summary of binding specificity (created in BioRender).

We then evaluated biomarker binding to IgG from other species including sheep and rabbit (**Figure 2C, D, and E**). Sheep-IgG was of particular interest due to the similar protein sequence with goat-IgG.^38^ Linear regression analysis indicates a correlation for sheep anti-cyanine IgG (1.69 ± 0.18; R^2^ = 0.75) and a weaker correlation for rabbit anti-RFP IgG (1.02 ± 0.27; R^2^ = 0.34) (**Figure 2**). The values for both the uncoated wells and rabbit anti-RFP IgG are significantly different from each other and from the values obtained when sheep- and goat-IgGs are used as capture antibodies (*p*<0.0001). The difference between median values and *p* value for each comparison are shown in **Figure S3C**. These data were then validated using competition ELISA by preincubating the human plasma with IgG derived from phylogenetically related (goat, sheep, cow) and unrelated species (mouse, rabbit) (**Figure S5**). Notably, antibodies from phylogenetically related species, but not IgG from unrelated species (*p*<0.0001), successfully compete with goat antibodies immobilized on the assay plate (**Figure 2F**). Taken together these results indicate the biomarker of interest has affinity for IgG from phylogenetically related species (goat, sheep, and cow), but not unrelated species (mouse or rabbit) (**Figure 2G**).

To determine if the biomarker is human-IgG, protein G affinity purification was used to isolate IgG from whole plasma. IgG concentrations were quantified in whole plasma (14.76 mg/mL) and compared to concentrations in retained (10.74 mg/mL) and non-retained (0.35 mg/mL; *p*<0.0001) fractions after protein G purification (**Figure 3A**). ELISA was then performed on the isolated fraction using a goat capture antibody resulting in similar biomarker levels in both whole plasma (median A450 = 0.45) and the IgG retained fraction (median A450 = 0.65), while the non-retained fraction approaches background levels (A450 = 0.15; *p*=0.01 relative to whole plasma; p<0.0001 relative to retained fraction) (**Figure 3B**). As confirmation, Coomassie stained SDS-PAGE gel of the isolated fractions show the heavy (∼55 kDa) and light (∼25 kDa) chains of IgG and no other co-eluted proteins (**Figure S6**). These results suggest that the biomarker observed in human plasma is likely a human-IgG targeting IgG from the *Bovidae* family.

**Figure 3.**
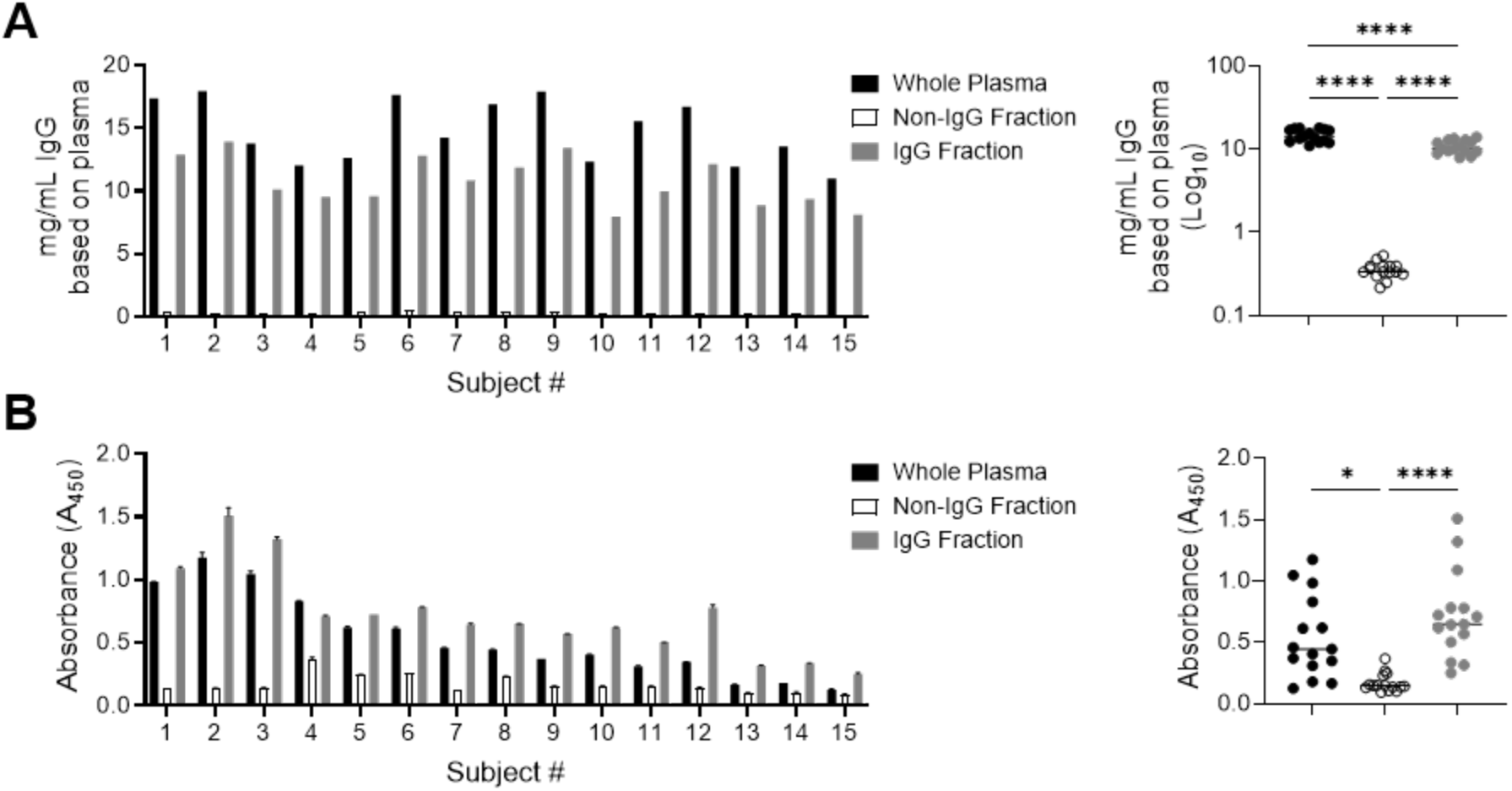
Antibody concentration and response to goat capture antibody after Protein G purification. **A**. IgG concentration and **B**. absorbance by ELISA before protein G (whole plasma) after protein G (Non-IgG Fraction) and after elution from protein G (IgG Fraction). **A**. IgG concentration calculated from standard curve using 4 parameter regression. One way ANOVA with Bonferroni’s multiple comparisons. Values in **B** are reported as median ± range. Friedman’s test with Dunn’s multiple comparisons. ** (*p*<0.01), *** (*p*<0.001) and **** (*p*<0.0001).

We then evaluated binding of the antibody biomarker to denatured goat-IgG by western blot using 15 plasma samples (**Figure S7**). These data indicate that the biomarker binds solely to the heavy chain of goat IgG (**Figure 4A**). The amount of denatured IgG on the membrane was increased and blotting repeated with plasma from subjects 1, 2, 3, 4, and 6 (**Figure 4B**), and complemented by competition ELISA with goat antibody fragments (Fc and Fab) (**Figure S8**). Whole antibody goat-IgG exhibits 38 ± 6% (mean ± SEM) interference as compared to -1 ± 6% for whole rabbit IgG. While the polyclonal nature of the biomarker causes variability in the competitive assay, whole goat-IgG is not statistically different from goat-Fc (51 ± 6%), and goat-Fab fragments (17 ± 5%) are statistically lower compared to whole goat-IgG (*p* = 0.03) and goat-Fc (*p* = 0.003). Collectively, the western blot and competition ELISA data indicate that antibody biomarker predominantly binds to the Fc portion of the goat-IgG heavy chain, and are not based on differences between human and goat IgG glycosylation patterns (**Figure S9**).^39^

**Figure 4.**
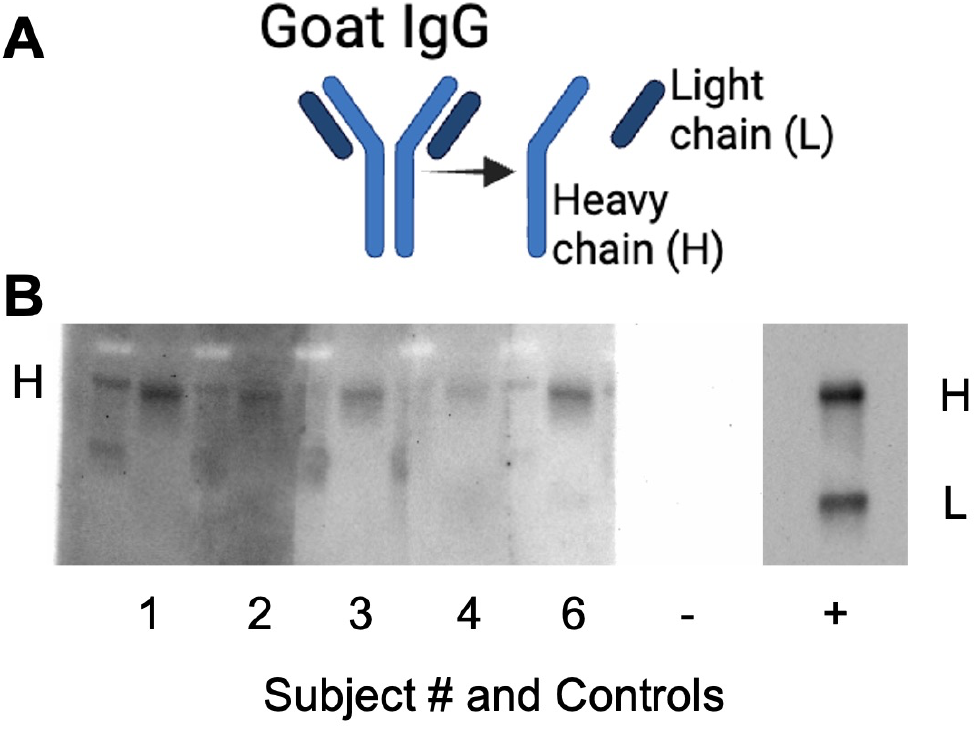
Identification of antibody biomarker binding target in goat-IgG by western blot. **A**. Summary of fragments from goat IgG evaluated by western blot including 25 kDa light chain (L) and 50 kDa heavy chain (H). **B**. Membranes with goat IgG probed with human plasma from 5 subjects who are biomarker positive by ELISA assays and exhibited binding in the preliminary western blot. Positive control from the same membrane was imaged separately (created with BioRender).

We then evaluated binding of the biomarker to goat-IgG in its native conformation using bio-layer interferometry (BLItz assay). Protein G purified IgG collected from subjects 1, 3, 5, and 15 were evaluated at multiple dilutions relative to whole plasma. A single dilution of 1:1.3 is plotted as a comparison between subjects using both anti-ApoA-I IgG and anti-GFP IgG (**Figure 5A**). A larger wavelength shift in this assay corresponds to increased binding to the target protein on the probe. Importantly, the wavelength shift magnitude for each sample aligns with the trend observed by ELISA (**Figure 5A**). This is consistent when using anti-ApoA-I and anti-GFP as different capture antibodies in this assay. Concentration dependent biomarker binding in replicates of plasma sample to both capture antibodies are shown in **Figure 5B** providing evidence of reproducible binding in multiple assays. The heterogeneity and low dissociation rate (*kd*) for these samples preclude affinity determination using this assay. Nonetheless, global fitting of these samples indicates binding affinity <10^-7^ M. Importantly, a lack of binding to purified polyclonal human-IgG (**Figure S10**) indicate that the antibody biomarker does not cross-react with human-IgG.

**Figure 5.**
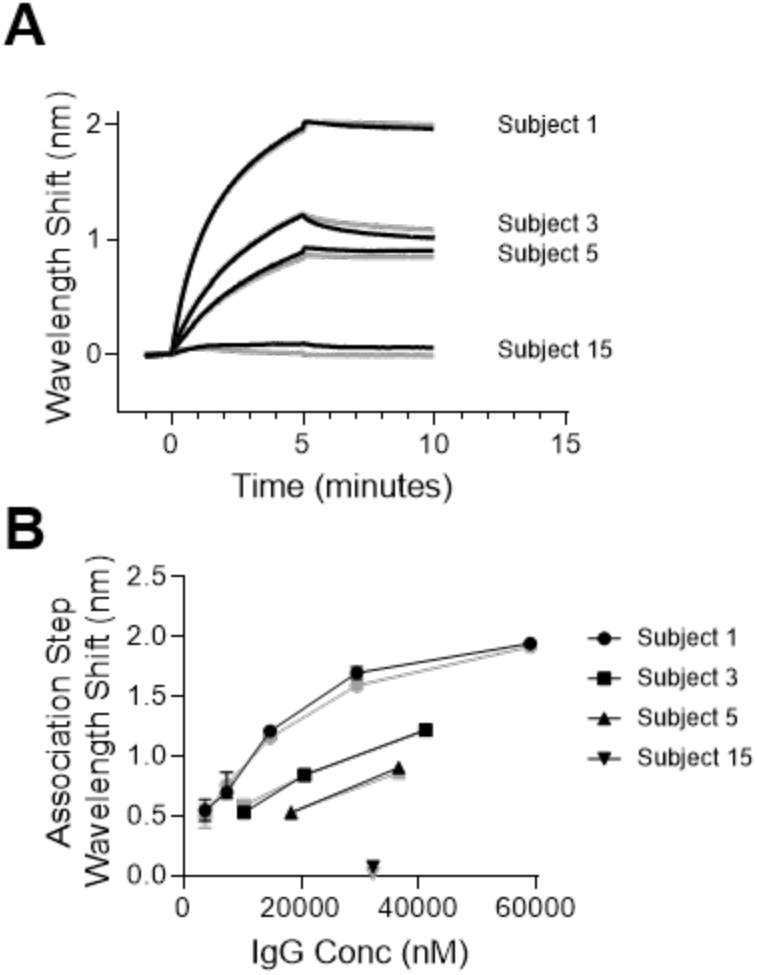
Binding kinetics of biomarker determined by bio-layer interferometry. **A**. Binding studies were completed with Protein G purified IgG fraction from four individual subjects (1, 3, 5, and 15) spanning the range of ELISA absorbance values. B. Maximum wavelength shift during the association step at multiple dilutions of Protein G purified IgG for the 4 subjects in A. Data from two independent experiments are presented with the median and range plotted. Samples in black are anti-ApoA-I capture antibody, and gray are anti-GFP capture antibody.

### Biomarker Validation in an Independent Cohort

To validate the antibody biomarker profiles in an independent cohort, we measured biomarker levels (Mean: 0.39 ± 0.38; IQR: 0.22-0.39) in 4356 baseline samples from the Multi-Ethnic Study of Atherosclerosis (MESA) as a prospective cohort (**Figure 6A**). The demographics and clinical characteristics of the analyzed samples are shown in **Table S3** and bivariate analysis between biomarker and characteristics shown in **Figure S11**. Initially, Cox regression analysis using low vs. high tertiles was conducted at four years of follow-up with any cardiovascular event (cvda, **Table S4**) as primary endpoint (**Figure S12**), and using covariates associated with three different models (**Table S5**). Notably, biomarker was significantly predictive up to 1-year after blood draw (unadjusted HR: 3.01 [95% CI, 1.28 – 7.08; *p*=0.012] between the lowest and the highest tertile; **Figure S12C, and D**), but not beyond 1 year, and remained significant after adjusting for covariates included in the Emory cohort (Model 1 adjusted HR: 2.77 [95% CI, 1.17 – 6.57; *p*=0.021] between lowest and highest tertiles; **Figure S12C**) or covariates used in the PREVENT equation (Model 2 adjusted HR: 2.55 [95% CI, 1.07 – 6.09; *p*=0.035] between lowest and highest tertiles; **Figure S12C**).

**Figure 6.**
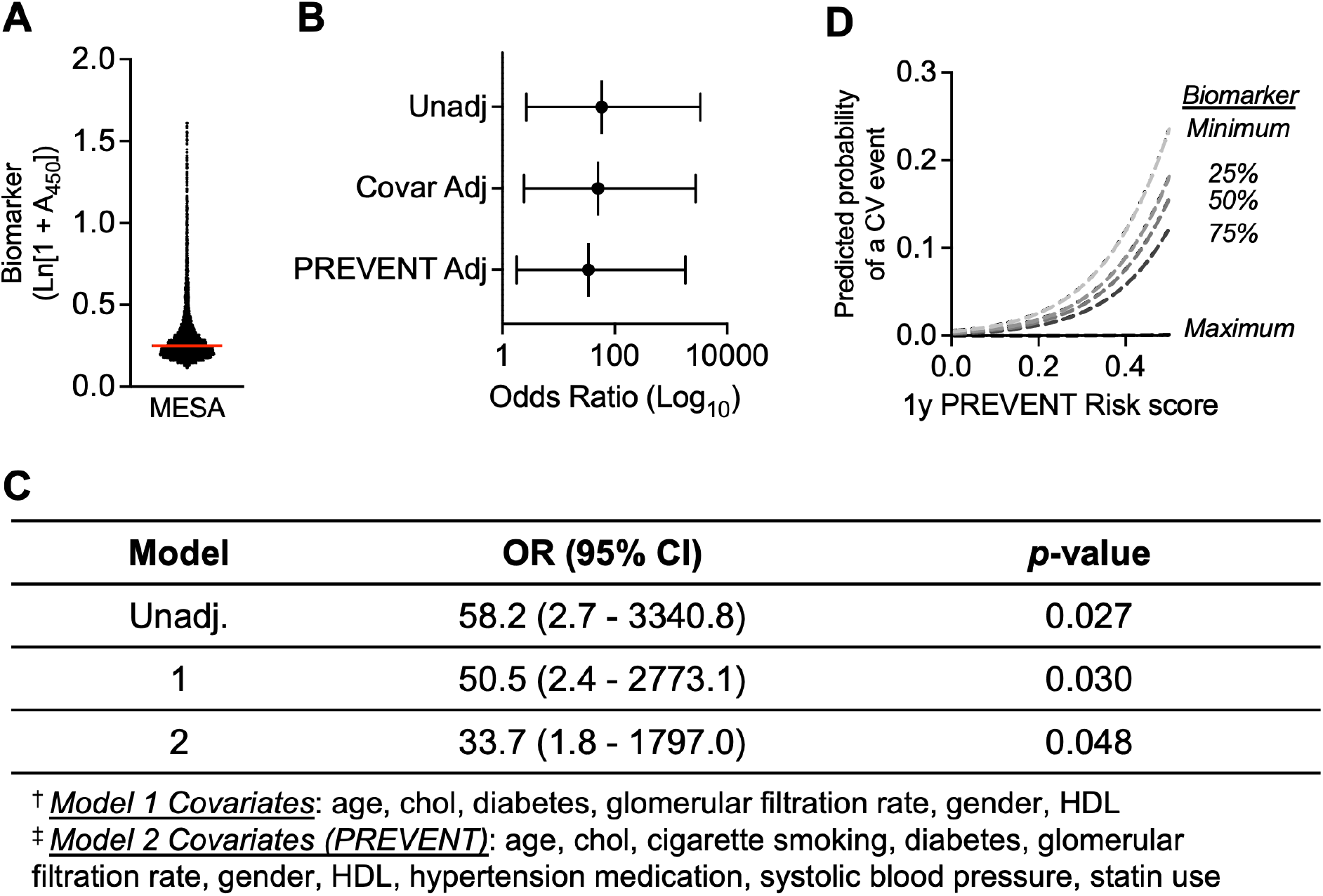
Logistic Regression analysis of patients from the MESA cohort. **A**. Biomarker levels were evaluated by ELISA at a plasma dilution of 1:200 in 4356 baseline samples from the prospective MESA cohort. LOG-transformed absorbance values are plotted, with median denoted by line. **B**. Odds ratio for unadjusted, covariate adjusted (Model 1) and PREVENT covariate adjusted outcomes at one year after blood collection. **C.** Proportional regression models used to determine odds ratios and CIs using covariates used in the CAD population in Figure 1 and a more sensitive PREVENT adjusted analysis. **D**. Biomarker levels stratify subjects at 1-year when used as a covariate in the PREVENT risk calculation, particularly in those at highest calculated risk of an event in which subjects at maximum biomarker levels have minimal relative risk.

Given the size of the cohort analyzed, biomarker level was also analyzed by logistic regression as a continuous variable at endpoints of 1-, 2-, 3-, 4-, and 18.5-years of follow-up. The primary endpoint was any cardiovascular event (cvda, **Table S4**) and secondary endpoints were independent analyses of non-fatal stroke, non-fatal MI, coronary heart disease, and cardiovascular death. By logistic regression, the unadjusted odds ratio was significant at one year after blood draw based on a 1-unit decrease in log-transformed biomarker (OR: 58.2; 95% CI: 2.7 – 3340.8; *p*=0.027) (**Figure 6B, C**), but not beyond one year (**Table S6**). Among the samples tested, there are 41 adjudicated endpoint events within one year of baseline blood draw. Although biomarker correlated with some variables (i.e. race/ethnicity, cholesterol, systolic blood pressure, diabetes mellitus, and smoking status) by bivariate analysis (**Figure S11**), the odds ratio remains significant after adjustment for covariates used in the original cohort (OR: 50.5; 95% CI: 2.4 – 2773.1; *p*=0.030) and covariates used in the PREVENT equation (OR: 33.7; 95% CI: 1.8 – 1797.0; *p*=0.048) (**Figure 6C**). Odds ratios were calculated using both the PREVENT and PCE risk scoring and adjusted for either the risk score alone as a covariate, or each individual variable included as a multivariate analysis. Notably, minimal differences are observed across each of the methods (**Table S6**).

Additionally, there was no correlation between biomarker levels and coronary artery calcification (**Figure S13**), suggesting that this biomarker provides incremental value for ASCVD risk prediction beyond that of coronary artery calcification testing. The biomarker was then used for risk stratification modeled at one year in combination with 10-year PREVENT risk prediction resulting in enhanced risk prediction when biomarker levels are included in the model, particularly in the population at highest risk (**Figure 6D**). Subjects at highest PREVENT-predicted risk and minimal biomarker levels have a 24.7% predicted chance of an event, while those with maximum biomarker levels have a 0.14% predicted chance of an event (**Figure 6D**).

## DISCUSSION

In this report we describe the use of an ELISA to measure an antibody biomarker which is inversely correlated with CVD events in patients with a history of CAD, and an independent prospective cohort of patients with no known clinical CVD. The observed association of this biomarker appeared independent of traditional risk factors and provided additional risk stratification beyond the former guideline-recommended PCE for estimating 10-year ASCVD risk^37^ and updated guideline-recommendation using the PREVENT equation.^36^ While patients with known CAD exhibit biomarker prediction over 4 years, patients without known clinical ASCVD exhibit an association between biomarker and incident ASCVD limited to a single year following sample collection.

Reasons for this observation are the topic of further study, including the temporal fluctuation of this novel biomarker in subjects over time.

Although, the assay was designed to measure ICs,^26, 29^ continued evaluation indicate that the biomarker is not related to ApoA-I, as previously described.^19^ Competition ELISA and protein G purification indicate that the biomarker is a human-IgG targeting antibodies from the *Bovidae* family identified serendipitously due to assay interference. While previous publications have identified human anti-animal antibodies, these reports focus on their role in interfering with clinical assays. To our knowledge this is the first report which finds an association between an anti-animal antibody profile and clinical outcomes.

Human anti-animal antibodies (HAAA) have been reported and are defined as highly specific antibodies, with strong binding to their target epitopes, and are thought to be the result of either iatrogenic and non-iatrogenic exposure.^11^ Alternatively, heterophilic antibodies are characterized by broad specificity and weak binding.^12^ Our data indicates that this biomarker more closely resembles a HAAA as we find that the biomarker is specific for IgG from the *Bovidae* family with no affinity for human-IgG. While biochemical characterization suggests that an exposure event may be the impetus for biomarker induction, the mechanisms leading to the antibody profile and the implications for cardiovascular health remain unclear.

The affinity of the antibody biomarker indicate that specific epitopes are likely involved in their induction, leading to somatic hypermutation and affinity maturation to achieve high affinity antibodies. The structural characteristics of IgG provide unique components to investigate antibody reactivity.^40^ Although humans do not produce N-glycolylneuraminic acid (NGNA) found on goat IgG,^41^ the biomarker binds de- glycosylated goat antibodies,^42^ which reduces the likelihood of this as an explanation for biomarker induction. More convincingly, western blot indicates binding to the IgG heavy chain, which is similar to work by Koshida and colleagues who found human anti-mouse antibodies that bind exclusively to the heavy chain of mouse antibodies.^43^

One possible explanation for the induction of antibodies targeting *Bovidae* IgG is dietary exposure to cow’s milk, which is known to induce an antibody response in infants, but these antibodies decrease over time due to oral tolerance.^44, 45^ Despite the induction of oral tolerance, Savilahta and colleagues detected anti-cow’s milk antibodies in control populations, including children from 7 to 14 years of age, an indication that milk-induced antibodies are present beyond early childhood.^46^ However, these studies do not specifically investigate antibody responses to cow-IgG.^47^ The phylogenetic similarities between IgG from species within the *Bovidae* family, including cow, may explain the high prevalence of the antibody biomarker in plasma samples collected from within the United States given the relatively high rates of milk and beef consumption, which could lead to a cross-reactive antibody response with goat-IgG.

An alternative explanation includes induction of cross-reactive antibodies to an unrelated pathogenic or symbiotic microbial protein through molecular mimicry. Molecular mimicry has been investigated in the context of autoimmune disorders, such as Epstein-Barr Virus and multiple sclerosis, in which an infection is implicated as the causative agent for induction of autoimmune pathology.^48^ Molecular mimicry between bacterial cell walls and oxidized LDL (oxLDL) is also an explanation for the induction of anti-oxLDL antibodies in CVD.^49^ Based on the outcome data observed with our assay, if the biomarker is induced due to molecular mimicry, the offending microorganism might be responsible for cardiac and/or vascular inflammation. Therefore, subjects who have an elevated, protective antibody biomarker to the microbe, and a corresponding cross-reactive response to goat-IgG could have reduced inflammation and improved outcomes. Several pathogens have been linked to atherosclerotic cardiovascular disease including *C. pneumoniae*, *P. gingivalis*, *H. pylori*, influenza A, hepatitis C, cytomegalovirus, and human immunodeficiency virus,^50^ but none have known protein sequence similarity with goat-IgG.

The paucity of known microbial proteins that exhibit sequence similarity with goat-IgG hinders antigen identification. However, structural motifs within goat-IgG may provide clues regarding the potential for antibody induction. Antibodies are part of the Ig superfamily (IgSF) group of proteins, which encompass a family of over 700 proteins expressed in the human genome.^51^ Additionally, Ig-like domains are ubiquitous in nature and have been characterized in many prokaryotic and eukaryotic organisms with limited sequence homology between species.^52^ Ig-like domains are characterized by their three-dimensional structural characteristic β-sandwich composed of anti-parallel β- strands, which result in increased stability and reduced protease susceptibility, making them ideal proteins as extracellular receptors or cell adhesion molecules in various species.^51^ An antibody response to a microbial protein with an Ig-like domain could minimize the effect of exposure and lead to cross reactivity with Ig-like domains from other species, including goat-IgG. Two common pathogens that express proteins with characterized Ig-like domains are Herpes virus and adenovirus.^53^ Importantly, 90% of the world’s population has one or both infections and much of the exposure occurs during childhood.^54, 55^ Other pathogenic or symbiotic microbes, including those found in the microbiome, with uncharacterized proteins containing Ig-like domains may also be responsible for antibody biomarker induction.^56^ Although the antigen responsible for induction of the biomarker remains unknown, the inverse correlation with CVD events prompts continued investigation.

The serendipitous discovery of a biomarker that correlates with cardiovascular outcomes is intriguing, but there are limitations. First, this report does not identify the target antigen responsible for induction of the biomarker and the target (goat-IgG) used in the assay may not be the actual target, which increases the potential for assay variability. Second, the temporal dynamics of biomarker levels in circulation are unknown and the temporal difference in predictive power between the CAD and non-CAD populations are unclear. Third, proper comparisons between each cohort used are not feasible given the differences in cohort characteristics, sample size, and respective outcomes. Finally, the sensitivity and specificity of the biomarker assay are necessary for clinical utility, but are outside the scope of this study. The limitations listed are the focus of ongoing research to build a more robust understanding of the biomarker, the antigen responsible, and the clinical translational potential.

In conclusion, antibodies that bind goat-IgG are present in human plasma and are inversely associated with CVD events in both a high-risk population and a prospective cohort. The biomarker possesses high affinity interaction with goat-IgG due to either direct exposure to the target or to an unrelated protein through molecular mimicry. Our findings have broad implications in clinical and biomedical sciences as a potential prognostic indicator and target for mechanistic and therapeutic studies to reduce the burden of CVD in patients. These data encourage continued evaluation of the antibody characteristics, functional implications and mechanisms leading to biomarker induction.

## ACKNOWLEDGMENTS

We thank the Kentucky Blood Center and Dr. Abdel-Latif for plasma samples. We also think Dr. Alan Daugherty for his valuable feedback and access to western blot instrumentation in his laboratory. We also thank the UK Proteomics core for use of the BLItz instrument. Images were prepared using BioRender.

## ABBREVIATIONS

ApoA-I: Apolipoprotein A-I
ApoB: Apolipoprotein B
ApoA-I/IgG ICs: ApoA-I/IgG immune complexes
ASCVD: Atherosclerotic cardiovascular disease
BLItz: Bio-layer interferometry
BME: beta-mercaptoethanol
CAD: Coronary artery disease
CBAL: Central blood analysis laboratory
CV: Coefficient of variation
CVD: Cardiovascular disease
ECL: Enhanced chemiluminescence
EDTA: Ethylenediaminetetraacetic acid
ELISA: Enzyme linked immunosorbent assay
GFP: Green fluorescent protein
HDL: High density lipoprotein
HAAA: Human anti-animal antibody
HRP: Horseradish peroxidase
ICs: Immune complexes
IgG: Immunoglobulin G
IQR: Interquartile range
KBC: Kentucky blood center
LDL: Low density lipoprotein
MESA: Multi-Ethnic Study of Atherosclerosis
MI: Myocardial infarction
NGNA: N-glycoylneuraminic acid
OxLDL: oxidized low density lipoprotein
PBS-C: Phosphate buffered saline with 0.1% casein and 0.1% tween
PBS-T: Phosphate buffered saline with 0.1% tween
PCE: Pooled cohort equation
PREVENT: Predicting Risk of cardiovascular EVENTs equation
TMB: Tetramethylbenzidine
UACR: Urine albumin-to-creatinine ratio

## HIGHLIGHTS

- Human IgG antibodies reactive with goat-IgG are inversely correlated with adverse cardiovascular events in both high and low risk subjects.
- The antibody biomarker is independent of common cardiovascular risk factors.
- The antibody biomarker is a high affinity antibody profile that bind the heavy chain of goat-IgG and IgG from other species in the *Bovidae* family.

