## Supplemental Information for "Identification and characterization of a human antibody profile inversely associated with adverse cardiovascular events"

***Short title***: Antibody profile associated with CVD events

**Corresponding Author:**
Vincent J. Venditto, PhD
University of Kentucky
Department of Pharmaceutical Sciences
Lee T. Todd Jr., Building, Room 337
789 S Limestone St.
Lexington, KY 40536-0596


#### MATERIALS AND METHODS

##### Human samples

**Patients With CAD:** Obstructive CAD was defined as visible plaque resulting in at least 50% luminal stenosis in at least 1 epicardial vessel. Those with a history of coronary artery bypass grafting or percutaneous coronary intervention were all labeled as having obstructive CAD. Estimated glomerular filtration rate was calculated using the Chronic Kidney Disease Epidemiology Collaboration equation. Heart failure was defined by the presence of self-reported history of heart failure or physician diagnosis of heart failure noted in medical record irrespective of ejection fraction. History of MI was diagnosed according to the American College of Cardiology/American Heart Association guidelines as self-reported chest pain and evidence of cardiac ischemia on 12-lead electrocardiogram or elevated cardiac enzymes greater than twice the upper limit of normal with angiographic findings consistent with MI diagnosis.

Patients enrolled in the Emory Cardiovascular Biobank, a prospective cohort of patients undergoing left heart catheterization for suspected or confirmed CAD at 3 Emory Healthcare sites in Atlanta, GA, were followed for determination of major adverse cardiovascular events (death/MI/stroke). Participants were interviewed to collect demographic characteristics, medical history, medication use, and behavioral habits. Risk-factor prevalence was determined by physician diagnosis and treatment for hypertension, hyperlipidemia, and diabetes mellitus, and these factors were also defined according to the Joint National Committee, Adult Treatment Panel III, and American Diabetes Association criteria,[1-3] respectively, and smoking habits were recorded. History of significant CAD was defined as at least 1 major epicardial vessel with ≥50% stenosis. Medical records and International Classification of Diseases, Ninth Revision (ICD-9) diagnostic codes were reviewed to confirm self-reported medical history. Cause of death was adjudicated by 2 independent cardiologists blinded to the data with a third arbitrator in case of disagreement. MI was diagnosed according to the American College of Cardiology/American Heart Association guidelines as presence of self-reported chest pain and evidence of cardiac ischemia on 12-lead electrocardiogram or elevated cardiac enzymes greater than twice the upper limit of normal with angiographic findings consistent with acute MI diagnosis. Outcome data were obtained by phone contact, electronic medical record review, and data from the social security death index and state records. Adjudication was conducted by personnel blinded to the data. The study was approved by the institutional review board at Emory University (Atlanta, GA). All subjects provided written informed consent at the time of enrollment.

##### Biomarker ELISA

**General Protocol**: Streptavidin coated pre-blocked plates (Thermofisher, 15124) were washed 3 times with 200 µL of phosphate buffered saline with 0.1% Tween (PBS-T). Capture antibody was immobilized using 100 µL of PBS-T containing 500 fmol (72 ng) of biotinylated goat anti-ApoA-I IgG (Abcam, ab27630, Lot #: GR171443-19) added to each well. The plates were incubated for 2h at 37 °C. Plates were washed 6 times with 200 µL of PBS-T prior to the addition of 100 µL of plasma at a final dilution of 1:200 in PBS with 0.1% casein (PBS-C). Following the addition of plasma, plates were incubated for 30 min at 37 °C. After incubation, plates were washed 6 times with PBS-T, then 100 µL of a 1:4000 dilution of horseradish peroxidase (HRP) conjugated secondary antibody (goat anti-human-IgG (HRP) (Abcam, ab7153)) were added to each well. Plates were incubated for either 30 min at 37 °C for Figures 1, S1 and S2 or 1h at 37 °C for the remaining figures before washing 6 times with 200 µL PBS-T. The ELISA was then developed with 100 µL of room temperature tetramethylbenzidine (TMB) incubated for 30 min at room temperature in the dark. The reaction was quenched with 100 µL of 0.5 M $H_2SO_4$ and resultant absorbance was measured at 450 nm (BioTek, Synergy Hybrid Reader or BMG ClarioStar). Samples were evaluated in duplicate and analyzed for coefficient of variation (CV) to confirm low replicate variability. Samples with CV > 10% were reanalyzed. ELISA validation assays are shown in **Figure S1 and S2**.

Competition ELISA were performed as described for standard ELISA with a final solution of 1:200 dilution of plasma and a ratio of 500:1, 10:1 or 0.2:1 of added IgG to capture IgG immobilized on the plate.

**Coefficients of Variation**: Wells of the streptavidin plates were either coated with biotinylated goat anti-ApoA-I IgG or left uncoated. Plasma from 18 blood donor subjects was diluted 1:200 with PBS-C and tested in 12 coated wells and 12 uncoated wells on an individual day. This was repeated until all 18 subjects had been tested 3 times. Results from these replicates are shown in **Figure S1**. Wells from subject 1 which had absorbance values over the instrument maximum were graphed as 4.0, and due to the frequent maximum values observed, subject 1 was excluded from any coefficient of variation analysis (CV). CV was calculated by taking the standard deviation of samples and dividing it by the average of the samples, for intra-assay CV the 12 values on a single day are utilized, for inter-assay CV all 36 values are used in the calculation. The average intra-assay CV was found to be 9.7% and the average inter-assay CV was determined to be 14.1%.

**Assay Linearity**: The linearity of the assay was evaluated in 12 samples at dilutions ranging from 1:50 to 1:400 and compared with capture-antibody-free wells at each plasma dilution (**Figure S2A**). One sample exhibited very high absorbance across all dilutions and was further diluted down to 1:12,800 (**Figure S2A)**. Given the differing concentrations of plasma at each dilution, a second linearity assay with constant plasma concentration of a 1:200 was completed by mixing diluted plasma from subjects with high responses and low responses at ratios of 4:0, 3:1, 2:2, 1:3, and 0:4 (**Figure S2B**).

**Alternative capture antibody ELISA:** Biomarker ELISA procedure was completed as indicated previously with use of alternative capture antibodies to coat the plate after initial washing. The antibodies utilized include: goat anti-ApoA-I IgG (Abcam, ab27630), goat anti-ApoB IgG (Abcam, ab20898), goat anti-GFP IgG (Abcam, ab6658), sheep anti-Cyanine IgG (Abcam, ab7625) and rabbit anti-RFP IgG (Abcam, ab34771). Alternative capture antibody ELISA and all MESA samples were washed using BioTek EL406 plate washer, all other ELISA were washed by hand using multi-channel pipet.

**Alternative secondary antibody ELISA:** Biomarker ELISA procedure was completed as indicated previously with use of alternative secondary antibodies. The antibodies utilized include: Goat anti-human IgG (HRP) (Abcam, ab7153), goat anti-Fluorescein IgG (HRP) (Abcam, ab6656), and Rabbit anti-human IgG (HRP) (Abcam, ab7160).

**Competition ELISA**

Competition ELISA were performed as described for standard ELISAs, with free antibodies mixed with plasma prior to incubation with assay plates. IgG used for competition was purchased from Rockland Immunochemicals (Goat IgG 005-0102-0010, Rabbit IgG 011-0102-0010, Mouse IgG 010-0102-0005, Sheep IgG 013-0102-0010, Cow IgG 001-0102-0010) and for use in this assay is diluted to 0.72 µg/µL in PBS-C, and serially diluted 1:50 with PBS-C. Each dilution of IgG is mixed 1:1 with 1:100 dilution of plasma in PBS-C. This final solution has 1:200 dilution of plasma and a ratio of 500:1, 10:1 or 0.2:1 of added IgG to capture IgG immobilized on the plate. As control, 1:100 dilution of plasma is also mixed 1:1 with PBS-C for a final dilution of 1:200 of human plasma and tested with or without initial coating via capture antibody. A commercial rabbit anti-goat-IgG (Abcam, ab97105) was tested along with the human plasma samples at a final dilution of 1:20,000). Percent interference was calculated based on the following equation:

$$\text{percent interference} = \frac{\text{No interference} - \text{Interference sample}}{\text{No interference} - \text{non specific binding}} \times 100$$

#### Protein G Spin Purification

Protein G spin columns (Thermofisher, 89979) were used based on the manufacturer's instructions. Columns were centrifuged at 1000 xg for 1 minute after each wash step or to collect samples. To prepare the column they were washed 2 times with 2 mL of wash buffer. Next, 2 mL of a 1:10 dilution of plasma in wash buffer was added to the column and mixed end over end for 10 minutes at room temperature. Flow through (non-retained fraction) was collected from the column and saved for ELISA. The column was then washed 3 times with 2 mL of wash buffer. To collect the bound fraction, 1 mL of elution buffer was added to the column and collected into a new 15 mL tube with 150 µL of neutralization buffer. This elution step was repeated 2 times with collection into the same 15 mL tube for a final total volume of 3.15 mL. Columns were regenerated by washing with 3 mL of elution buffer, 3 mL of wash buffer and 2 additional washes with 2 mL of wash buffer.

During this procedure the non-IgG fraction and whole plasma controls were diluted to 1:10 with wash buffer as a part of this procedure. For the elution, a total of 200 µL of plasma was added to the column in the initial step and 3150 µL of total elution volume were collected for an approximate final dilution of 1:16.

To prepare samples for the Bio-Layer Interferometry (BLItz) assay this procedure was repeated and the IgG fraction was concentrated via Protein concentrator tubes with a 10 kDa molecular weight cutoff (Thermofisher, 88516).

#### IgG Quantification ELISA

High binding plates (Microlon 600) were washed 3 times with 200 µL of $Na_2CO_3$ buffer (9.7 pH, 0.05 M). Donkey anti-human-IgG (Abcam, ab102421) was diluted 1:500 in $Na_2CO_3$ buffer and added to the plates. Plates were incubated 1 hour at 37 °C and then washed 6 times with 200 µL of PBS-T. Plates were then blocked with 200 µL of PBS-C for 1 hour at 37 °C. The control and IgG fraction samples were diluted to a final dilution of 1:4,000,000 while the non-IgG fraction was adjusted to a final dilution of 1:50,000. Following a wash with PBS-T, samples were transferred to the assay plate. Control samples with known concentration were added to each plate to generate an 8-point standard curve with human-IgG (Genscript, A01006). Assay plates with samples were incubated for 30 minutes at 37 °C and then washed 6 times with 200 µL of PBS-T and a 1:4000 dilution of goat anti-human-IgG (HRP) (Abcam, ab7153) was added to each well. These plates were incubated for 1 hour at 37 °C. Plates were washed a final 6 times with 200 µL of PBS-T and then 100 µL of room temperature TMB was added to each plate and incubated in the dark for 30 minutes at room temperature. The reaction was quenched with 100 µL of 0.5 M $H_2SO_4$ and the resultant absorbance was measured at 450 nm (BioTek, Synergy Hybrid Reader). Samples were evaluated in duplicate. IgG concentration was calculated based on four parameter regression based on the standard curve on each plate.

**Western Blots**

Goat-IgG (Rockland Immunochemical, 005-0102-0010) was diluted to 90 µg/mL in PBS then mixed with 4x buffer containing beta-mercaptoethanol (BME) and heated at 95 °C for 5 min. Mini-PROTEAN TGX Precast Protein Gel (4-20%; Biorad, 4561096) were loaded with either 6.67 µL of denatured goat-IgG or 2.5 µL of PageRuler Plus Prestained Protein Ladder (Thermofisher, 26619) in alternating lanes with 7 samples and 8 ladders per gel. Gels were electrophoresed on ice at 100 V. Transfers were completed using TurboBlot system with Trans-Blot Turbo Mini 0.2 µm PVDF Transfer Pack (Biorad, 1704156). Membranes were blocked with Superblock PBS (Thermofisher, 37515) overnight at 4 °C. Membrane sections containing IgG and ladder were cut from the membrane to stain individually with human plasma diluted 1:400 in PBS-C with 0.1% Tween (PBS-CT) for 1h at RT. Membranes were washed with PBS-T and incubated with anti-human-IgG secondary antibody (Abcam, ab7153) at a dilution of 1:10,000 in PBS-CT for 1h at RT. Positive controls were evaluated on membrane sections without plasma by incubating with a 1:40,000 dilution of rabbit anti-goat-IgG (Abcam, ab6741). Negative controls were evaluated by incubating the membrane sections with PBS-CT lacking human plasma, followed by incubation with anti-human-IgG secondary antibody. Membranes were imaged with enhanced chemiluminescence (ECL) (ThermoFisher, 32209) and figures prepared with Biorad image processing software (Biorad, ChemiDoc MP). Samples that bound goat-IgG by western blot were reanalyzed using a 120 µg/mL solution of goat-IgG in PBS.

**Deglycosylation of Goat-IgG**

Deglycosylated antibodies were prepared based on a modified protocol from Kaneko et al using PNGaseF.[4] PNGaseF (500 units) (New England Biolabs, P0704S) was mixed with 2 mg of goat-IgG and incubated at 37 °C for 41 hours. A protein G spin column was then used to collect the IgG from the reaction mixture according to the procedure described in the methods section. Collected IgG was compared to untreated IgG and IgG treated with PNGaseF under denaturing conditions via SDS-PAGE.

**SDS-PAGE of deglycosylated Goat-IgG and Protein G purified plasma samples**

IgG samples were denatured in 4x loading buffer with BME and placed in an electrophoresis chamber set at 100 V in a 10% mini-PROTEAN TGX Precast Protein Gels (Biorad, 4561036) and stained with coomassie blue to label proteins. For deglycosylated samples, the migration of heavy chain in each condition was compared to confirm deglycosylation was complete and IgG from each reaction condition was utilized in a competition ELISA at 100 times the quantity of capture antibody, untreated goat-IgG and rabbit-IgG were used as controls. For protein G purified samples, 4-20% gels were run at 150 V and assessed for bands other than IgG heavy and light chains after concentration with Protein concentrator tubes with a 10k molecular weight (MW) cutoff (Thermofisher, 88516).

**Bio-Layer Interferometry (BLItz) Assay**

Binding kinetics of the biomarker were measured by Bio-Layer Interferometry, (BLItz Octet N1, Sartorius). Baseline interferometry for a streptavidin biosensor (Sartorius, 18-5020) was collected in Octet Kinetics buffer (Sartorius, 18-1105) and the sensor was then coated with biotinylated goat-IgG at 12.5 µg/mL for 3 min. After the coating step, a baseline was established for 3 min in kinetics buffer. Association between biomarker and goat-IgG was measured by incubating the probe with samples containing biomarker for 5 min. Biomarker dissociation was measured by placing the probe in kinetics buffer for 5 min. At each concentration, the samples were evaluated with an uncoated probe, as well as a probe coated with anti-ApoA-I IgG (Abcam, ab27630) and a probe coated with biotinylated goat anti-GFP IgG (Abcam, ab6658). Data from the uncoated probe was subtracted from the assay experiment for data visualization. Experiments were repeated on two different days for reproducibility. BLItz Pro 1.3 software was used to collect data and determine kinetic constants using step correction and global fitting across multiple concentrations for anti-ApoA-I and anti-GFP. To assess binding to human IgG, the same experimental design was performed with biotinylated-human-IgG (Rockland, 009-0602) to coat the probe.

#### SUPPLEMENTAL TABLES

**Table S1**. Demographics and clinical characteristics of patients from Emory Cardiovascular Biobank with CAD.

| **Characteristic** | **Data (n = 359)** |
|---|---|
| Age, median, yr (SD) | 64.2 (13.1) |
| Male sex, n (%) | 231 (64.7) |
| Black race, n (%) | 84 (23.5) |
| Body mass index, kg/m$^2$, median, (SD) | 29.2 (5.7) |
| Systolic blood pressure, mmHg (SD) | 138 (20) |
| Diastolic blood pressure, mmHg (SD) | 71 (20) |
| Hypertension, n (%) | 314 (88) |
| Hyperlipidemia, n (%) | 274 (76.8) |
| Diabetes mellitus, n (%) | 120 (33.6) |
| Obstructive CAD (>=50%), n (%) | 267 (74.8) |
| Previous myocardial infarction, n (%) | 84 (23.6) |
| Acute myocardial infarction, n (%) | 42 (11.8) |
| Heart failure, n (%) | 109 (30.5) |
| Ejection fraction, n (%) | 53 (14) |
| Total cholesterol, mg/dL, median (SD) | 158 (47) |
| Low density lipoprotein, mg/dL, median (SD) | 92 (43) |
| High density lipoprotein, mg/dL, median (SD) | 41 (15) |
| Creatinine, mg/dL, median (IQR) | 1.5 (1.9) |

**Table S2**. Spearman correlation between biomarker levels and baseline characteristics in subjects from the Emory Cardiovascular Biobank.

| **Baseline Characteristic (n=359)** | **Biomarker level Rho (*p*-value)** |
|---|---|
| Male | 0.07 (0.22) |
| Black | 0.00 (0.98) |
| Age | -0.01 (0.84) |
| Body mass index | -0.03 (0.59) |
| Systolic blood pressure | -0.02 (0.75) |
| Diastolic blood pressure | 0.03 (0.56) |
| Acute myocardial infarction | 0.02 (0.74) |
| Obstructive coronary artery disease | -0.05 (0.31) |
| History of myocardial infarction | -0.03 (0.53) |
| History of heart failure | 0.00 (0.95) |
| Diabetes | -0.09 (0.10) |
| Hypertension | -0.09 (0.10) |
| Hyperlipidemia | 0.02 (0.71) |
| Total cholesterol | 0.05 (0.32) |
| Triglycerides | 0.00 (0.99) |
| High density Lipoprotein | -0.06 (0.23) |
| Low density lipoprotein | 0.07 (0.19) |
| Ejection Fraction | 0.07 (0.20) |
| Creatinine | -0.04 (0.43) |
| Glomerular Filtration Rate | 0.08 (0.13) |
| *hs-CRP (n = 193) | -0.07 (0.30) |

**Table S3**. Demographics and clinical characteristics of analyzed MESA samples.

| Characteristic | Data (n = 4356) |
|---|---|
| Age, median, yr (SD) | 61 (9.9) |
| Male sex, n (%) | 1934 (44.4) |
| White subjects, n (%) | 1776 (40.8) |
| Chinese subjects, n (%) | 508 (11.7) |
| Black subjects, n (%) | 1145 (26.3) |
| Hispanic subjects, n (%) | 927 (21.3) |
| Body mass index, $kg/m^2$, median, (SD) | 28.4 (5.6) |
| Systolic blood pressure, mmHg (SD) | 126 (21) |
| Diastolic blood pressure, mmHg (SD) | 72 (10) |
| Hypertension, n (%) | 1873 (43.0) |
| Diabetes mellitus, n (%) | 523 (12.0) |
| Total cholesterol, mg/dL, median (SD) | 196 (36) |
| Low density lipoprotein, mg/dL, median (SD) | 118 (32) |
| High density lipoprotein, mg/dL, median (SD) | 51 (15) |
| Creatinine, mg/dL, median (IQR) | 0.92 (0.30) |

**Table S4.** Data entry codes used as variables in analysis of MESA cohort samples.

| Variable | Description | Criteria (if any) | Value Labels |
|---|---|---|---|
| agatpm1c | Mean: Agatston calcium score, phantom-adjusted | | |
| age1c | Age | | |
| bmi1c | Body mass index (kg/m^2) | | |
| chda | Coronary heart disease (CHD), all | | 0 = No<br>1 = Yes |
| chf | Congestive heart failure (CHF) | | 0 = No<br>1 = Yes |
| chol1 | Total cholesterol (mg/dL) | | |
| cig1c | Cigarette smoking status | | 0 = Never<br>1 = Former<br>2 = Current |
| cvda | Cardiovascular Disease (CVD), all | MI<br>Resusitated cardiac arrest<br>Definite Angina<br>Probable Angina (if followed by revascularization)<br>Stroke<br>Stroke Death<br>CHD Death<br>Other Atherosclerotic Death<br>Other CVD Death | 0 = No<br>1 = Yes |
| dm031c | Exam 1 Diabetes mellitus by 2003 ADA Fasting Criteria Algorithm | | 0 = Normal<br>1 = IFG<br>2 = Untreated diabetes<br>3 = Treated diabetes |
| dth | Death | | 0 = No<br>1 = Yes |
| dthtype | Death type | | 1 = Atherosclerotic coronary heart disease<br>2 = Stroke<br>3 = Atherosclerotic disease other than coronary disease,stroke<br>4 = Other cardiovascular disease, not defined above<br>5 = Non-cardiovascular disease<br>6 = Death type unkown, no death certificate<br>9 = Non-cardiovascular disease (ineligible for review)<br>999 = Pending adjudication |
| egfr1c | MDRD EGFR Based on Exam 1 Scale CR | | |
| gender1 | Gender | | 0 = Female<br>1 = Male |

| | | | |
|---|---|---|---|
| hdl1 | HDL cholesterol (mg/dL) | | |
| htnmed1c | Hypertension medication | | 0 = No<br>1 = Yes |
| mi | Myocardial infarction (mi) | | 0 = No<br>1 = Yes |
| race1c | Race / Ethnicity | | 1: White<br>2: Chinese<br>3: Black<br>4: Hispanic / Latino |
| sbp1c | Seated systolic blood pressure (mmHg) | | |
| strk | Stroke | | 0 = No<br>1 = Yes |
| sttn1c | HMG CoA Reductase inhibitors (statins) | | 0 = No<br>1 = Yes |
| ualbcre1 | Urinary albumin/creatinine (mg/g) | | |

**Table S5**. Variables and corresponding variable code used in covariate adjustment models in MESA cohort.

| Variable | Description | Model 1 | Model 2 (PREVENT) | Model 3 (PCE) |
|---|---|---|---|---|
| age1c | Age | X | X | X |
| bmi1c | BMI | | X | |
| chol1 | Cholesterol | X | X | X |
| cig1c | Cigarette smoking status | | X | X |
| dm031c | Diabetes mellitus | X | X | X |
| egfr1c | Estimated glomerular filtration rate | X | X | X |
| gender1 | Gender | X | X | X |
| hdl1 | HDL cholesterol | X | X | X |
| htnmed1c | Hypertenstion medication | | X | X |
| race1c | Race/ethnicity | | | X |
| sbp1c | Systolic blood pressure | | X | X |
| sttn1c | Statin use | | X | |

**Table S6**. Biomarker logistic regression analysis in baseline MESA samples using non-fatal stroke, non-fatal MI, and cardiovascular death as endpoints.

| Model | Timepoint (yrs) | Odds Ratio | 95% CI | p-value |
|---|---|---|---|---|
| Unadjusted | 1 | 58.17 | 2.65 – 3340.84 | 0.027 * |
| | 2 | 2.45 | 0.65 – 13.31 | 0.24 |
| | 3 | 3.11 | 0.98 – 12.82 | 0.08 |
| | 4 | 2.01 | 0.79 – 6.09 | 0.18 |
| | 18.5 | 0.75 | 0.51 – 1.14 | 0.17 |
| Model 1 covariate adjusted† | 1 | 50.47 | 2.43 – 2773.07 | 0.030 * |
| | 2 | 2.23 | 0.59 – 11.95 | 0.29 |
| | 3 | 2.91 | 0.91 – 12.03 | 0.10 |
| | 4 | 1.95 | 0.76 – 5.94 | 0.20 |
| | 18.5 | 0.73 | 0.49 – 1.12 | 0.15 |
| Model 2 PREVENT covariate adjusted‡ | 1 | 33.72 | 1.79 – 1796.99 | 0.048 * |
| | 2 | 2.02 | 0.56 – 10.51 | 0.34 |
| | 3 | 2.48 | 0.80 – 10.05 | 0.16 |
| | 4 | 1.70 | 0.68 – 5.05 | 0.30 |
| | 18.5 | 0.67 | 0.44 – 1.03 | 0.06 |
| Model 2.2 PREVENT score adjusted§ | 1 | 37.71 | 2.10 – 1863.37 | 0.037 * |
| | 2 | 2.36 | 0.67 – 11.91 | 0.24 |
| | 3 | 3.03 | 0.99 – 11.84 | 0.08 |
| | 4 | 2.01 | 0.81 – 5.89 | 0.17 |
| | 18.5 | 0.73 | 0.48 – 1.12 | 0.14 |
| Model 3 PCE covariate adjusted# | 1 | 36.95 | 1.95 – 1982.32 | 0.043 * |
| | 2 | 2.05 | 0.56 – 10.87 | 0.34 |
| | 3 | 2.72 | 0.87 – 10.92 | 0.12 |
| | 4 | 1.84 | 0.73 – 5.44 | 0.23 |
| | 18.5 | 0.70 | 0.46 – 1.08 | 0.10 |
| Model 3.2 PCE score adjusted§ | 1 | 45.61 | 2.26 – 2489.76 | 0.034 * |
| | 2 | 2.34 | 0.63 – 12.54 | 0.26 |
| | 3 | 3.02 | 0.95 – 12.42 | 0.09 |
| | 4 | 1.97 | 0.77 – 6.01 | 0.19 |
| | 18.5 | 0.69 | 0.46 – 1.06 | 0.06 |

† _Model 1 Covariates_: age, cholesterol, diabetes mellitus, estimated glomerular filtration rate, gender, HDL

‡ _Model 2 Covariates (PREVENT)_: age, cholesterol, cigarette smoking, diabetes mellitus, estimated glomerular filtration rate, gender, HDL, hypertension medication, systolic blood pressure, statin medication

### _Model 3 Covariates (PCE)_: age, cholesterol, cigarette smoking, diabetes mellitus, estimated glomerular filtration rate, gender, HDL, hypertension medication, race/ethnicity, systolic blood pressure

§ Score adjusted reflects an adjustment based on the 10-year risk predictive scoring using PREVENT and PCE equations.

#### SUPPLEMENTAL FIGURES

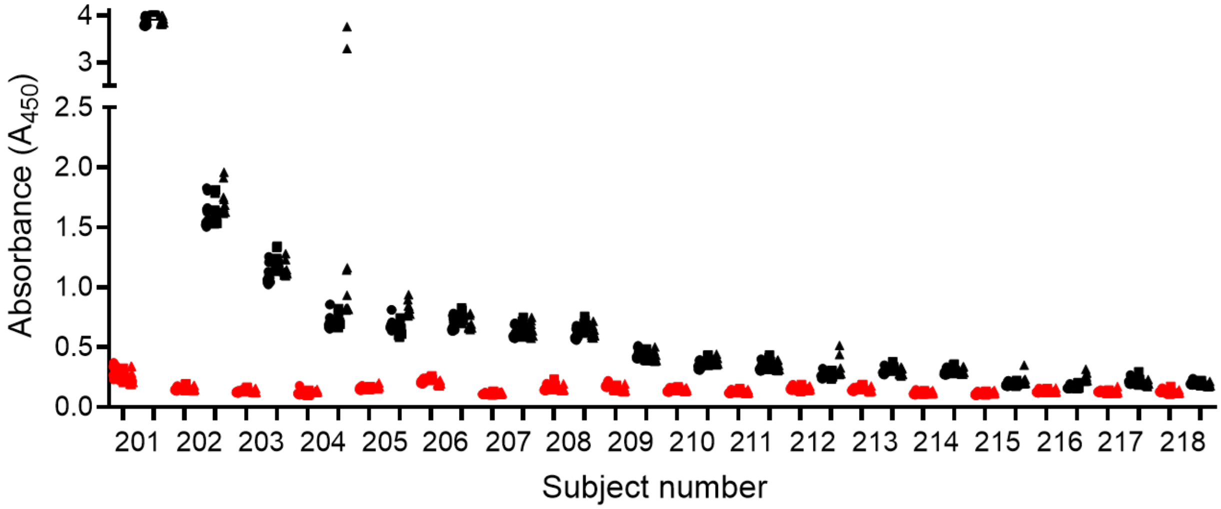


**Figure S1**. **ELISA validation assays to determine intra- and inter-assay coefficients of variation.** Plasma samples from 16 subjects from the Kentucky Blood Center were analyzed by ELISA in 12 coated (Black symbols) and 12 uncoated wells (red symbols) to determine intra-assay coefficients of variation. Samples were evaluated on three days to determine inter-assay coefficient of variation.

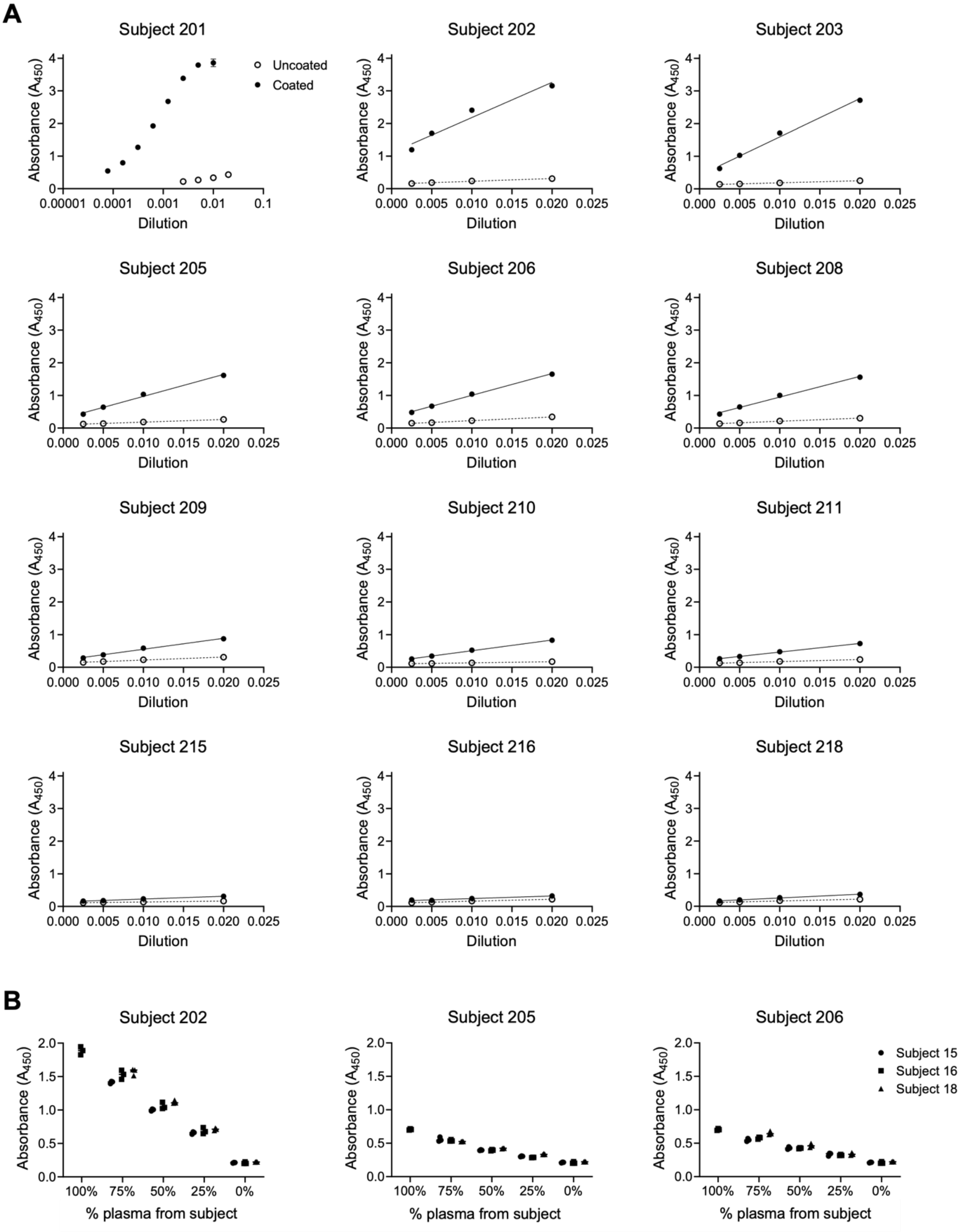

A
Subject 201
Subject 202
Subject 203
Subject 205
Subject 206
Subject 208
Subject 209
Subject 210
Subject 211
Subject 215
Subject 216
Subject 218
Absorbance ($A_{450}$)
Dilution
Uncoated
Coated
B
Subject 202
Subject 205
Subject 206
% plasma from subject
100%
75%
50%
25%
0%
Subject 15
Subject 16
Subject 18

**Figure S2**. **Validation of linearity of the assay. A**. ELISA was completed with dilutions ranging from 1:50 to 1:400 in 12 subjects. Subject 1 exhibited very high levels of the biomarker and was further diluted to 1:12,800 and is plotted on a log scale. Closed circles and solid line represent coated wells while open circles and dotted line represent uncoated wells. **B**. ELISA was completed at a constant 1:200 dilution of plasma where plasma from three high subjects was mixed with plasma from three low subjects (Subjects 15, 16, 18) at ratios of 4:0, 3:1, 2:2, 1:3 and 0:4.

**A**

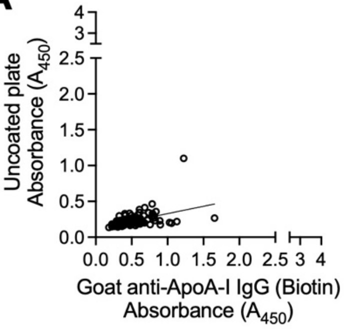


**B**

| Coating antibody | Slope ± 95% CI | $R^2$ |
|---|---|---|
| Uncoated | 0.20 ± 0.07 | 0.21 |
| Goat anti-ApoB IgG | 0.97 ± 0.13 | 0.65 |
| Goat anti-GFP IgG | 1.02 ± 0.08 | 0.86 |
| Sheep anti-cyanine IgG | 1.68 ± 0.18 | 0.75 |
| Rabbit anti-RFP IgG | 1.02 ± 0.27 | 0.34 |

**C**

| | Goat | | | Sheep | Rabbit |
|---|---|---|---|---|---|
| | anti-ApoA-I | anti-ApoB | anti-GFP | anti-Cyanine | anti-RFP |
| Uncoated | 0.27 **** | 0.18 **** | 0.24 **** | 0.28 **** | 0.66 **** |
| Goat anti-ApoA-I IgG | | 0.09 ** | 0.03 ns | 0.02 * | 0.39 **** |
| Goat anti-ApoB IgG | | | 0.07 *** | 0.11 **** | 0.48 **** |
| Goat anti-GFP IgG | | | | 0.04 ns | 0.42 **** |
| Sheep anti-Cyanine IgG | | | | | 0.37 **** |

**Figure S3. Effect of plate coating for biomarker ELISA. A**. Absorbance values using goat anti-ApoA-I IgG as the capture antibody in comparison to an uncoated plate. **B**. Linear regression analysis data from comparative ELISA assays. **C**. Absolute difference of median values and differences between groups determined via Friedman test with Dunn's multiple comparislon for absorbance values in Figure 2E. Not significant (ns), * ($p$<0.05), **($p$<0.01), ***($p$<0.001), ****($p$<0.0001).

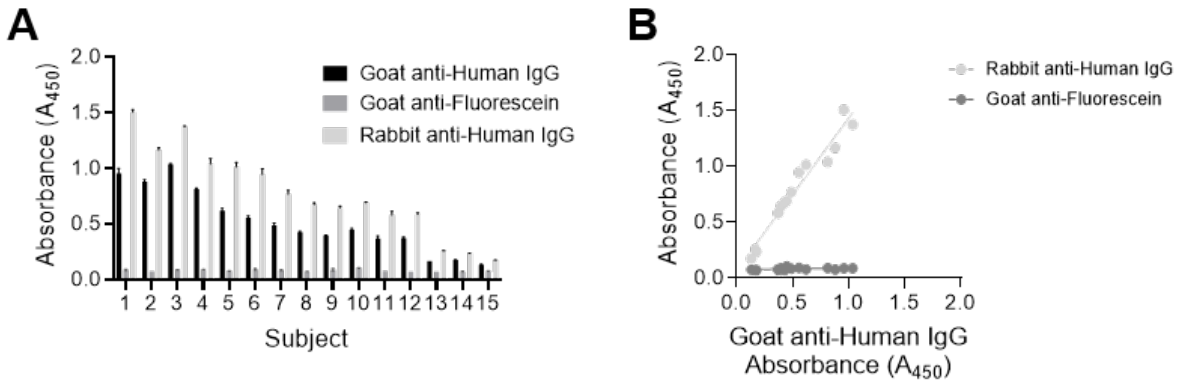


**Figure S4**. **Evaluation of secondary antibody effects in ELISA.**
**A**. Bar graph and **B**. Scatter plot using goat anti-ApoA-I IgG with (HRP)-conjugated goat anti-human IgG, (HRP)-conjugated goat anti-fluorescein and (HRP)-conjugated rabbit anti-human IgG as secondary antibodies in ELISA for 15 samples with known absorbance values. **A** Samples evaluated in duplicate with median and range presented.

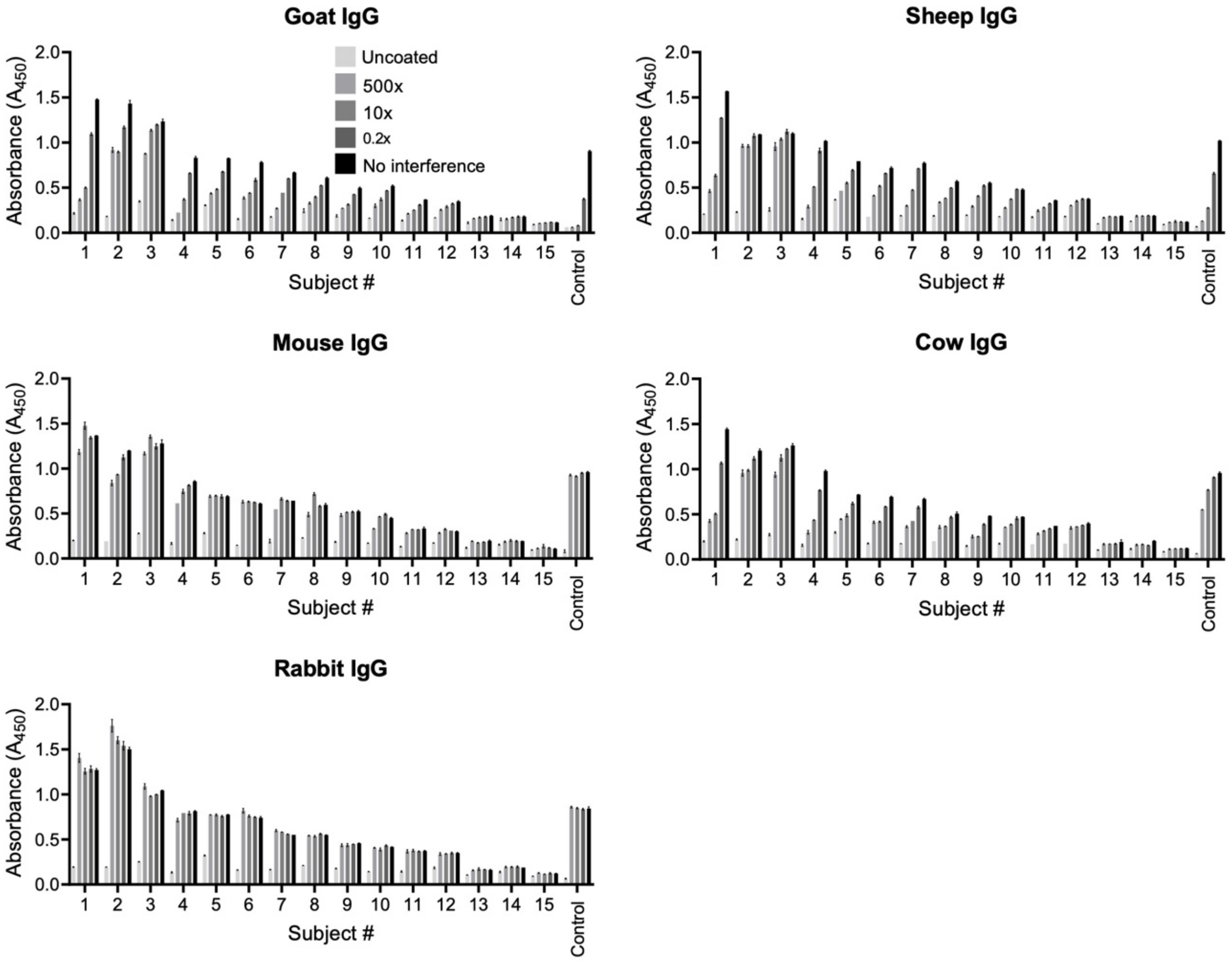


**Figure S5. Interference of biomarker measurement using control IgG competitors from different species.** Fifteen plasma samples with known antibody response, as determined by ELISA, evaluated in competition with goat-, mouse-, rabbit-, sheep- and cow-IgG. IgG from each species was mixed with plasma prior to addition to assay plate at different concentrations relative to capture antibody (500x, 10x, 0.2x and 0x). Rabbit anti-goat IgG is included as a control. Samples are run in duplicate and presented as median ± range.

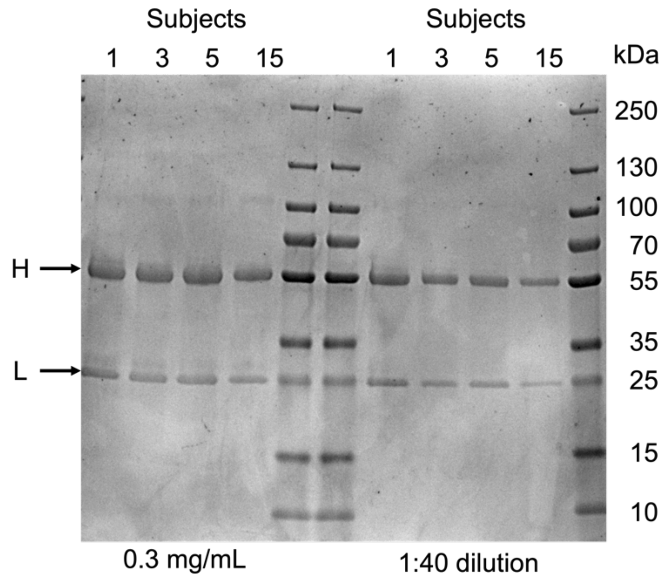


**Figure S6**. **SDS PAGE of Protein G purified plasma samples**.
Protein G purified plasma samples used for BLItz assay evaluated at constant concentration across samples (0.3 mg/mL) or constant dilution (1:40 based on whole plasma). Gel brightness was increased by 40% using Microsoft Word Picture Format Corrections to clearly show protein bands in this document.

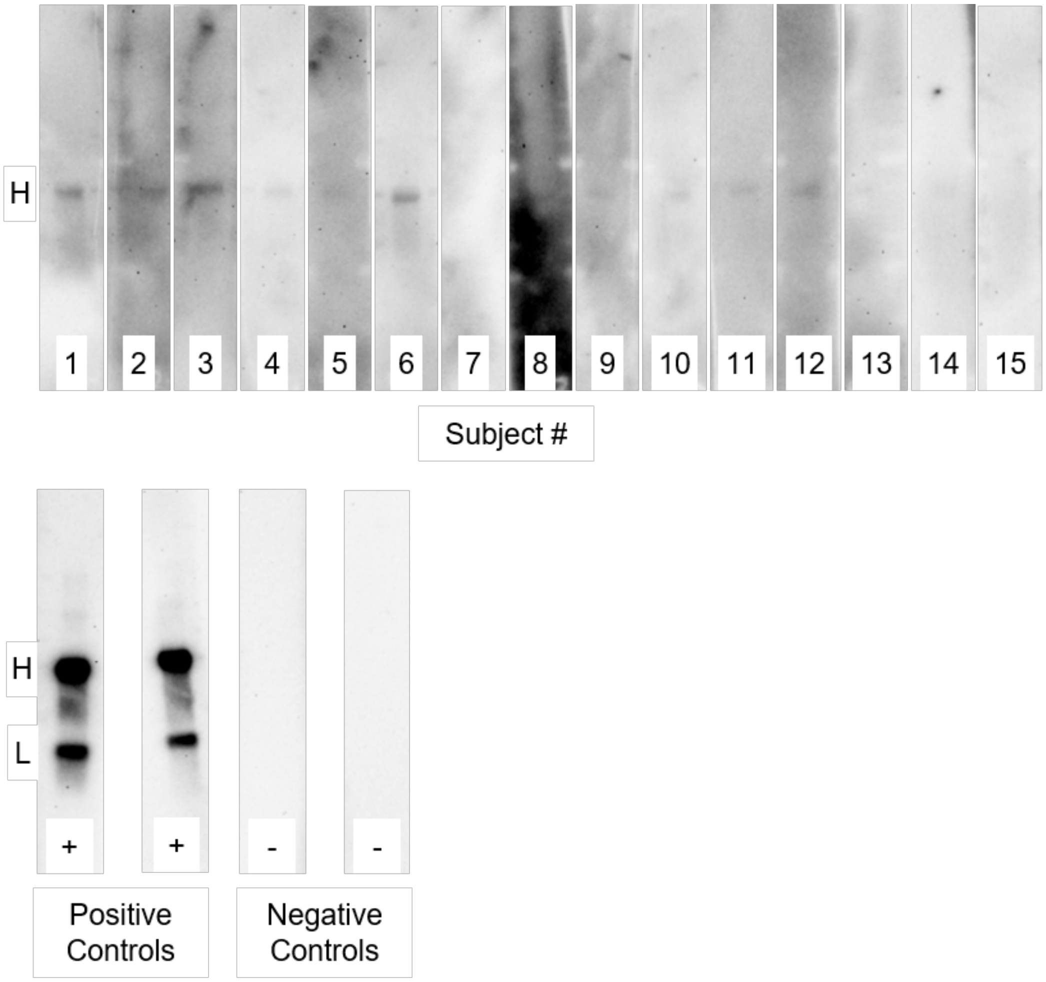


**Figure S7**. **Western blot of goat IgG using human plasma samples containing biomarker as source of primary antibody**.
Denatured goat-IgG was probed with human plasma from the 15 subjects along with positive controls utilizing a commercial anti-goat antibody and negative control with no plasma additions. "H" denotes approximately 50 kDa the size of the IgG heavy chain while "L" denotes approximately 25 kDa the size of the light chain.

**A**

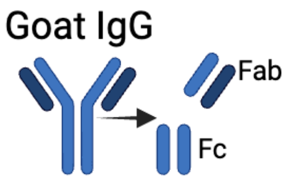


**B**

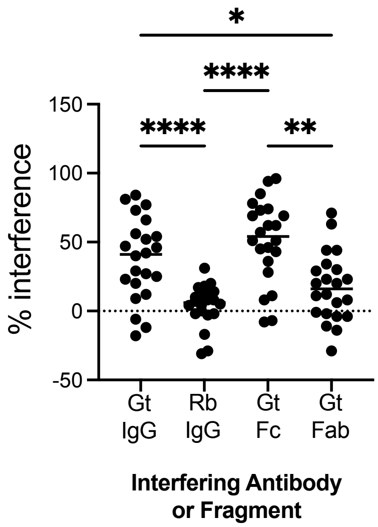


**Figure S8. Interference ELISA of biomarker samples with goat antibody fragments**.

Plasma samples from 22 blood donor subjects were evaluated by interference ELISA using whole goat IgG as capture antibody and either goat IgG, rabbit IgG, goat Fc, or goat Fab as interfering agents at 100x concentration relative to goat-IgG capture antibody. Data are evaluated by Friedman's test with Dunn's multiple comparisons. One sample in the Rb IgG condition resulted in -112% interference and is excluded from the graph for clarity, but included in statistical analysis.

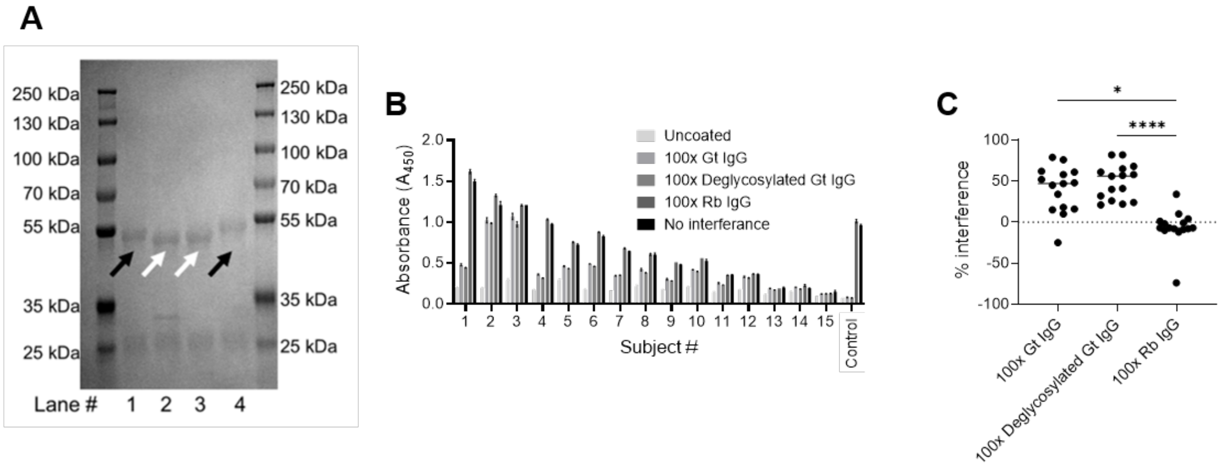


**Figure S9**. **Interference assay with deglycosylated goat IgG**.
**A**. Coomassie Stained gel. Lane 1, untreated goat IgG. Lane 2, control complete PNGase reaction. Lane 3, isolated PNGase treated goat IgG. Lane 4, untreated goat IgG. Black arrows point to heavy chain of untreated goat IgG. White arrows to the lower molecular weight deglycosylated IgG. **B**. A450 values and **C**. interference values for deglycosylated IgG and controls for **B** and **C**. **C** Friedman's test with Dunn's multiple comparison. ** ($p<0.01$), **** ($p<0.0001$).

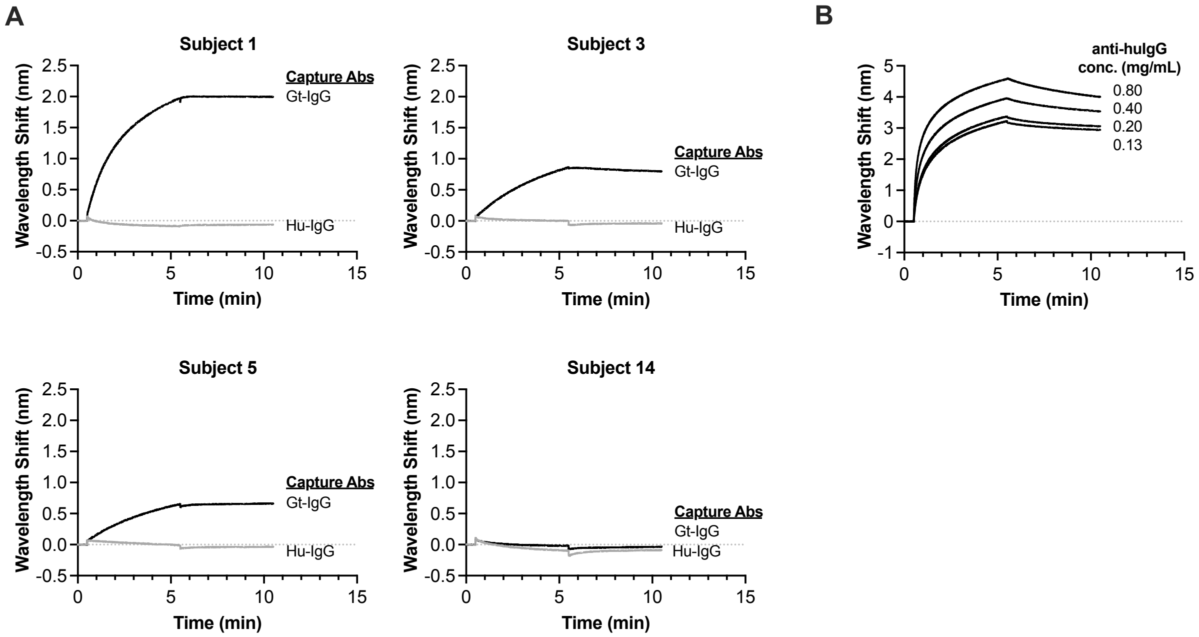


**Figure S10**. **Binding of purified biomarker to goat-IgG and human-IgG determined by bio-layer interferometry**. **A.** Binding studies were completed with Protein G purified IgG fraction from four inidividuals (1, 3, 5, and 14) spanning the range of biomarker ELISA absorbance values. Data from two independent experiments are presented with median plotted. Samples in black lines donate binding to goat-IgG, grey lines denote binding to human-IgG. **B.** Binding of control anti-human-IgG at four concentrations to human-IgG capture antibody by BLItz assay.

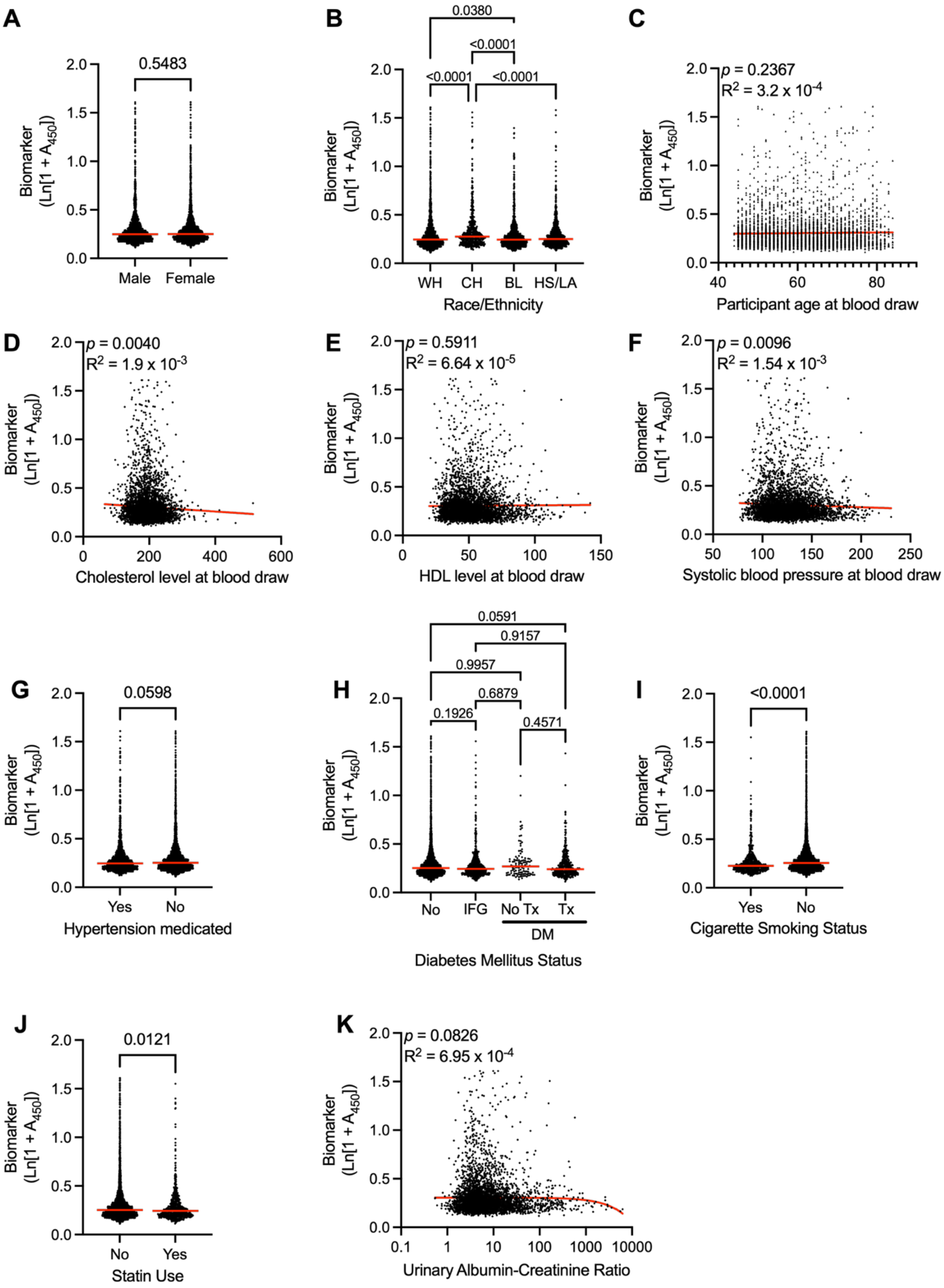
A
0.5483
Biomarker (Ln[1 + $A_{450}$])
Male
Female
B
0.0380
<0.0001
<0.0001
<0.0001
WH
CH
BL
HS/LA
Race/Ethnicity
C
$p$ = 0.2367
$R^2 = 3.2 \times 10^{-4}$
Participant age at blood draw
D
$p$ = 0.0040
$R^2 = 1.9 \times 10^{-3}$
Cholesterol level at blood draw
E
$p$ = 0.5911
$R^2 = 6.64 \times 10^{-5}$
HDL level at blood draw
F
$p$ = 0.0096
$R^2 = 1.54 \times 10^{-3}$
Systolic blood pressure at blood draw
G
0.0598
Yes
No
Hypertension medicated
H
0.0591
0.9157
0.9957
0.6879
0.1926
0.4571
No
IFG
No Tx
Tx
DM
Diabetes Mellitus Status
I
<0.0001
Yes
No
Cigarette Smoking Status
J
0.0121
No
Yes
Statin Use
K
$p$ = 0.0826
$R^2 = 6.95 \times 10^{-4}$
Urinary Albumin-Creatinine Ratio

**Figure S11. Bivariate analysis between biomarker and demographic and clinical data associated with MESA samples.** Variables shown are based on variables used in the PREVENT cardiovascular risk prediction calculation. **A**. Gender; **B**. Race/Ethnicity (WH: White; CH: Chinese; BL: Black; HS/LA: Hispanic/Latino; **C**. Age; **D**. Total cholesterol; **E**. HDL cholesterol; **F**. Systolic blood pressure; **G**. Medicated for hypertension; **H**. Diabetes Mellitus (IFG: impaired fasting glucose; No Tx: untreated; Tx: treated); **I**. Cigarette smoking status (N: never; F: former; C: current); **J**. Statin use; **K**. Urinary albumin-creatinine ratio (UACR).

**A**

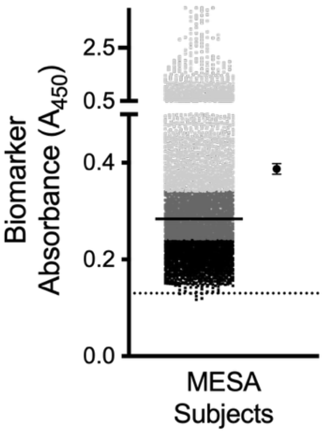


**B**

| Lowest vs. Highest Tertile | | HR (95% CI) | p-value |
|---|---|---|---|
| 1 year | Unadjusted | 3.01 (1.28 – 7.08) | **0.012 *** |
| | Adjusted (Model 1) | 2.77 (1.17 – 6.57) | **0.021 *** |
| | Adjusted (Model 2) | 2.55 (1.07 – 6.09) | **0.035 *** |
| | Adjusted (Model 3) | 2.54 (1.06 – 6.06) | **0.036 *** |
| 4 year | Unadjusted | 1.22 (0.84 – 1.78) | 0.294 |
| | Adjusted (Model 1) | 1.12 (0.77 – 1.64) | 0.547 |
| | Adjusted (Model 2) | 1.08 (0.74 – 1.58) | 0.699 |
| | Adjusted (Model 3) | 1.09 (0.75 – 1.60) | 0.651 |

*Model 1 variables*: age, gender, cholesterol, diabetes mellitus, estimated glomerular filtration rate, HDL.
*Model 2 variables (PREVENT)*: age, gender, race, cholesterol, diabetes mellitus, HDL, hypertension medicated, systolic blood pressure, cigarette smoking
*Model 3 variables (PCE)*: age, gender, race, cholesterol, diabetes mellitus, HDL, hypertension medicated, systolic blood pressure, cigarette smoking

**C**

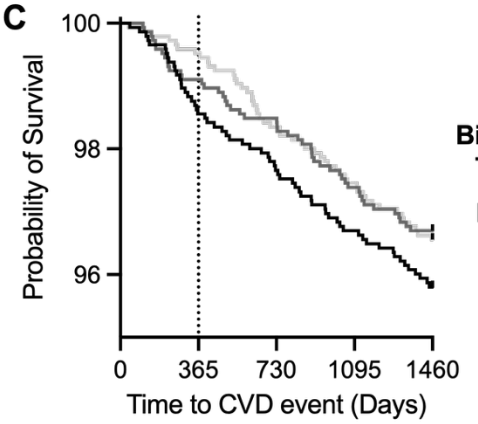


**D**

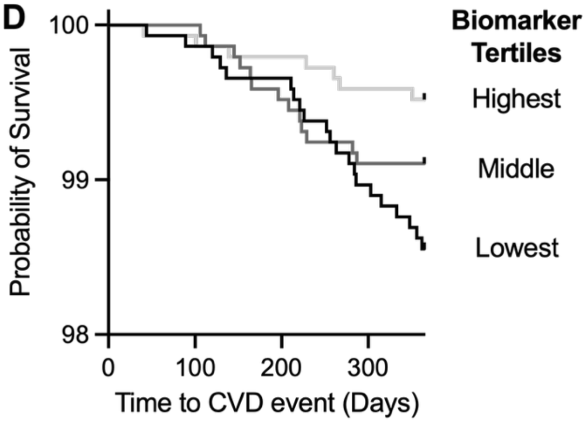


**Figure S12. Cox proportional regression analysis of MESA subjects based on biomarker levels. A.** Biomarker levels were evaluated by ELISA at a plasma dilution of 1:200 in 4356 baseline samples from the MESA cohort. Absorbance values are plotted, with median noted by line, tertiles are color coded to indicate highest (light gray), middle (dark gray), low (black). The mean value and 95% CI are plotted to the right. Dashed line denotes the limit of detection, defined as 3 SD above plasma-free wells. **B**. Proportional regression models used to determine hazard ratios using on established risk predictors of cardiovascular disease progression. **C, D**. Kaplan-Meier plot of major adverse cardiovascular events over 4 (**C**) and 1 (**D**) years of follow-up are shown. Subjects are divided into tertiles by absorbance in the biomarker assay as denoted in A.

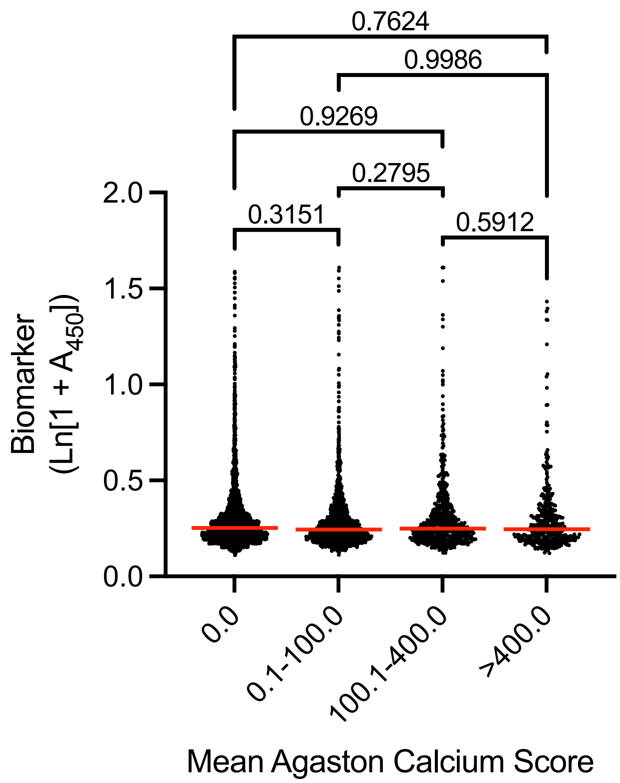


**Figure S13**. Biomarker level is not correlated with coronary artery calcification in MESA cohort. Biomarker levels shown for 4356 samples analyzed from MESA cohort and stratified based on Agaston coronary artery calcification scoring.

#### Major Resources Table

In order to allow validation and replication of experiments, all essential research materials listed in the Methods should be included in the Major Resources Table below. Authors are encouraged to use public repositories for protocols, data, code, and other materials and provide persistent identifiers and/or links to repositories when available. Authors may add or delete rows as needed.

**Antibodies**

| Target antigen | Vendor or Source | Catalog # | Working concentration |
|---|---|---|---|
| Goat anti-human-ApoA-I | Abcam | Ab27630 | 500 fmol |
| Goat anti-human-IgG | Abcam | Ab7153 | 1:4000 |
| Goat anti-human-ApoB | Abcam | Ab20898 | 500 fmol |
| Goat anti-GFP | Abcam | Ab6658 | 500 fmol |
| Sheep anti-Cyanine | Abcam | Ab7625 | 500 fmol |
| Rabbit anti-RFP | Abcam | Ab34771 | 500 fmol |
| Goat anti-fluorescein | Abcam | Ab6656 | 1:4000 |
| Rabbit anti-human-IgG | Abcam | Ab7160 | 1:4000 |
| Donkey anti-human-IgG | Abcam | Ab102421 | 1:500 |
| Human IgG | Genscript | A01006 | 0.03-64 mg/mL |
| Goat IgG | Rockland Immunochemicals | 005-0102-00010 | 0.72 µg/mL |
| Rabbit IgG | Rockland Immunochemicals | 011-0102-0010 | 0.72 µg/mL |
| Mouse IgG | Rockland Immunochemicals | 010-0102-0005 | 0.72 µg/mL |
| Sheep IgG | Rockland Immunochemicals | 013-0102-0010 | 0.72 µg/mL |
| Cow IgG | Rockland Immunochemicals | 001-0102-0010 | 0.72 µg/mL |
| Rabbit anti-goat-IgG | Abcam | Ab97105 | 1:20,000 |
| Human-IgG biotinylated | Rockland Immunochemicals | 009-0602 | 12.5 µg/mL |

(1) Lenfant, C.; Chobanian Aram, V.; Jones Daniel, W.; Roccella Edward, J. Seventh Report of the Joint National Committee on the Prevention, Detection, Evaluation, and Treatment of High Blood Pressure (JNC 7). *Hypertension* **2003**, *41* (6), 1178-1179. DOI: 10.1161/01.HYP.0000075790.33892.AE (acccessed 2021/04/01).
(2) Brown, M. S.; Ho, Y. K.; Goldstein, J. L. The cholesteryl ester cycle in macrophage foam cells. Continual hydrolysis and re-esterification of cytoplasmic cholesteryl esters. *Journal of Biological Chemistry* **1980**, *255* (19), 9344-9352. DOI: https://doi.org/10.1016/S0021-9258(19)70568-7.
(3) Genuth, S.; Alberti, K. G.; Bennett, P.; Buse, J.; Defronzo, R.; Kahn, R.; Kitzmiller, J.; Knowler, W. C.; Lebovitz, H.; Lernmark, A.; et al. Follow-up report on the diagnosis of diabetes mellitus. *Diabetes Care* **2003**, *26* (11), 3160-3167. DOI: 10.2337/diacare.26.11.3160 From NLM.
(4) Kaneko, Y.; Nimmerjahn, F.; Ravetch, J. V. Anti-Inflammatory Activity of Immunoglobulin G Resulting from Fc Sialylation. *Science* **2006**, *313* (5787), 670-673.
